# PEAK3 pseudokinase represents a pro-migratory and -invasive signalling scaffold

**DOI:** 10.1101/2021.02.17.431740

**Authors:** Jianmei Hou, Elizabeth V Nguyen, Minglyanna Surudoi, Michael J Roy, Onisha Patel, Isabelle S Lucet, Xiuquan Ma, Roger J Daly

**Author notes:** Co-senior authors (Xiuquan Ma, Roger J Daly).

## Abstract

The PEAK family of pseudokinases comprises PEAK1 and PEAK2 as well as the recently-identified PEAK3. PEAK1/2 play fundamental roles in regulating tyrosine kinase signal output and oncogenesis, while PEAK3 remains poorly-characterized. Here, we demonstrate that PEAK3 undergoes homotypic association as well as heterotypic interaction with PEAK1/2. PEAK3 also recruits ASAP1/2, Cbl and PYK2 and the adaptors Grb2 and CrkII, with binding dependent on PEAK3 dimerization. PEAK3 tyrosine phosphorylation on Y24 is also dependent on dimerization as well as Src family kinase activity, and interestingly, is decreased via PTPN12 in response to EGF treatment. Y24 phosphorylation is required for binding of Grb2 and ASAP1. Overexpression of PEAK3 in MDA-MB-231 breast cancer cells enhanced cell elongation and cell motility, while knockdown of endogenous PEAK3 decreased cell migration. In addition, overexpression of PEAK3 in PEAK1/2 compound knock-out MCF-10A breast epithelial cells enhanced acinar growth and invasion in 3D culture, with the latter phenotype dependent on PEAK3 tyrosine phosphorylation and binding of Grb2 and ASAP1. These findings characterize PEAK3 as an integral member of the PEAK family with scaffolding roles that promote cell proliferation, migration and invasion.

## Introduction

Approximately 10% of human protein kinases are classified as pseudokinases since they lack at least one of the conserved amino acid motifs important for catalytic function (Manning *et al*, 2002; Reiterer *et al*, 2014). Although most of them are predicted to be catalytically inactive they play important signalling roles by functioning as scaffolds, anchors, or allosteric regulators (Reiterer *et al.*, 2014). Three pseudokinases, PEAK1 (also known as SgK269), PEAK2 (SgK223/Pragmin) and the recently-identified PEAK3 (Chromosome 19 Open Reading Frame 35, C19orf35) form the PEAK family. These three proteins share highly conserved domain structures including a C-terminal pseudokinase (PsK) domain with adjacent N- and C-terminal α-helical regions (Patel *et al*, 2020). In addition, they all contain specific N-terminal extensions, which are long in PEAK1 (∼ 1,200 residues) and PEAK2 (∼900 residues), but much shorter in PEAK3 (∼ 100 residues) (Patel *et al.*, 2020).

Both PEAK1 and PEAK2 function as scaffolds, harbouring tyrosine phosphorylation sites that specifically recruit src homology (SH)2 and phosphotyrosine binding (PTB) domain-containing effectors to regulate cell proliferation and/or migration/invasion. PEAK1 is phosphorylated by activated Src family kinases (SFKs) downstream of the epidermal growth factor receptor (EGFR) and specific integrins and regulates the p130Cas-Crk-paxillin, Erk and Shc1 signaling pathways to promote cell migration and invasion (Croucher *et al*, 2013; Wang *et al*, 2010; Zheng *et al*, 2013). Phosphorylation of PEAK1 Y635 creates a binding site for the Grb2 SH2 domain and leads to Ras signalling pathway activation and aberrant growth of MCF-10A mammary epithelial cells in 3D (Croucher *et al.*, 2013). Also, binding of PEAK1 Y1188 to the Shc1 PTB domain enables Shc1 to switch from regulating Grb2-dependent mitogenic activity to PEAK1-orchestrated cell morphology and invasion (Zheng *et al.*, 2013). Another important function of PEAK1 is that it represents a regulator of epithelial-mesenchymal transition (EMT) (Croucher *et al.*, 2013). For example, in MCF-10A mammary epithelial cells, overexpression of PEAK1 results in these cells converting to an elongated, mesenchymal morphology, and stable knockdown of PEAK1 in MDA-MB-231 breast cancer cells leads to mesenchymal-epithelial transition (MET) (Croucher *et al.*, 2013). Several studies over the last decade have reported PEAK1 overexpression in multiple human malignancies including breast (Agajanian *et al*, 2015; Croucher *et al.*, 2013), colon (Huang *et al*, 2018; Wang *et al.*, 2010), lung (Ding *et al*, 2018) and pancreatic cancers (Kelber *et al*, 2012), suggesting an oncogenic role in cancer growth and progression. In the case of PEAK2, the EPIY_411_A motif serves as a docking site for the SH2 domain of the C-terminal Src kinase (Csk), highlighting a critical role for PEAK2 in controlling Csk localisation to regulate SFK activity (Safari *et al*, 2011). Similar to PEAK1, PEAK2 induces cell elongation and co‐expression of Csk and PEAK2 markedly induces cell scattering (Senda *et al*, 2016). PEAK2 is required for the invasion of colon carcinoma cells (Leroy *et al*, 2009) and PEAK2 expression is significantly increased in pancreatic cancer and associated with poor prognosis in non-small cell lung cancer (Kong *et al*, 2016; Tactacan *et al*, 2015).

PEAK1 and PEAK2 form homo- and hetero-dimers, this representing an important regulatory mechanism in their downstream signaling and function (Ha & Boggon, 2018; Liu *et al*, 2016; Patel *et al*, 2017). Homo- and heterotypic association of PEAK1 and PEAK2 is mediated by the highly-conserved split helical dimerization (SHED) domain, comprised of the helix immediately N-terminal to the PsK domain and 3 helices C-terminal to the PsK domain that form an “XL”-shaped helical bundle (Ha & Boggon, 2018; Liu *et al.*, 2016; Patel *et al.*, 2017). The ability of PEAK1 to enhance cell elongation and migration is dependent on the functionality of the SHED region. In addition, PEAK1 requires PEAK2 to efficiently promote cell migration and activate Stat3 in MCF-10A cells and PEAK1 bridges PEAK2 to Grb2 (Liu *et al.*, 2016). Therefore, homo- and heterotypic association of these scaffolds is critical to biological activity and acts as a mechanism to diversify signalling outputs (Liu *et al.*, 2016).

The third family member, C19orf35 or PEAK3, was identified by the Roche group in 2018 (Lecointre *et al*, 2018), although its signalling mechanism and function remain unclear. Recently, it was demonstrated via site-directed mutagenesis and co-immunoprecipitation studies that PEAK3 also homo-dimerizes via the highly conserved SHED domain, this being critical for CrkII recruitment (Lopez *et al*, 2019). In addition, while PEAK3 overexpression did not impact cell morphology, it prevented CrkII-dependent membrane ruffling, leading the authors to suggest that PEAK3 may represent a negative modulator of PEAK1/2 action (Lopez *et al.*, 2019).

In this study, we characterize PEAK3 signalling mechanisms and function in detail. The resulting data indicate that PEAK3 has a scaffolding function, a pro-migratory and -invasive role, and demonstrate important functional roles for tyrosine phosphorylation, dimerization and recruitment of specific effectors including Grb2 and ASAP1.

## Results

### PEAK3 undergoes heterotypic association with PEAK1 and PEAK2

PEAK3 exhibits a similar molecular structure to PEAK1 and PEAK2 (Fig 1A). They all share a conserved C-terminal PsK domain with adjacent N- and C-terminal α-helical regions forming the highly-conserved split helical dimerization (SHED) domain, and a N-terminal extension which is significantly shorter in PEAK3. PEAK1 and PEAK2 harbour a CrkII SH3 domain binding motif (PPPLPKK) which is conserved in PEAK3 (residues 56-62) (Lopez *et al.*, 2019). Additionally, PEAK3 contains a SH2 binding motif (YSNL, residues 24-27) at the N-terminus and another potential SH3 binding motif (PGAPWR, residues 243-248), located in the PsK domain (Fig 1A).

**Figure 1.**
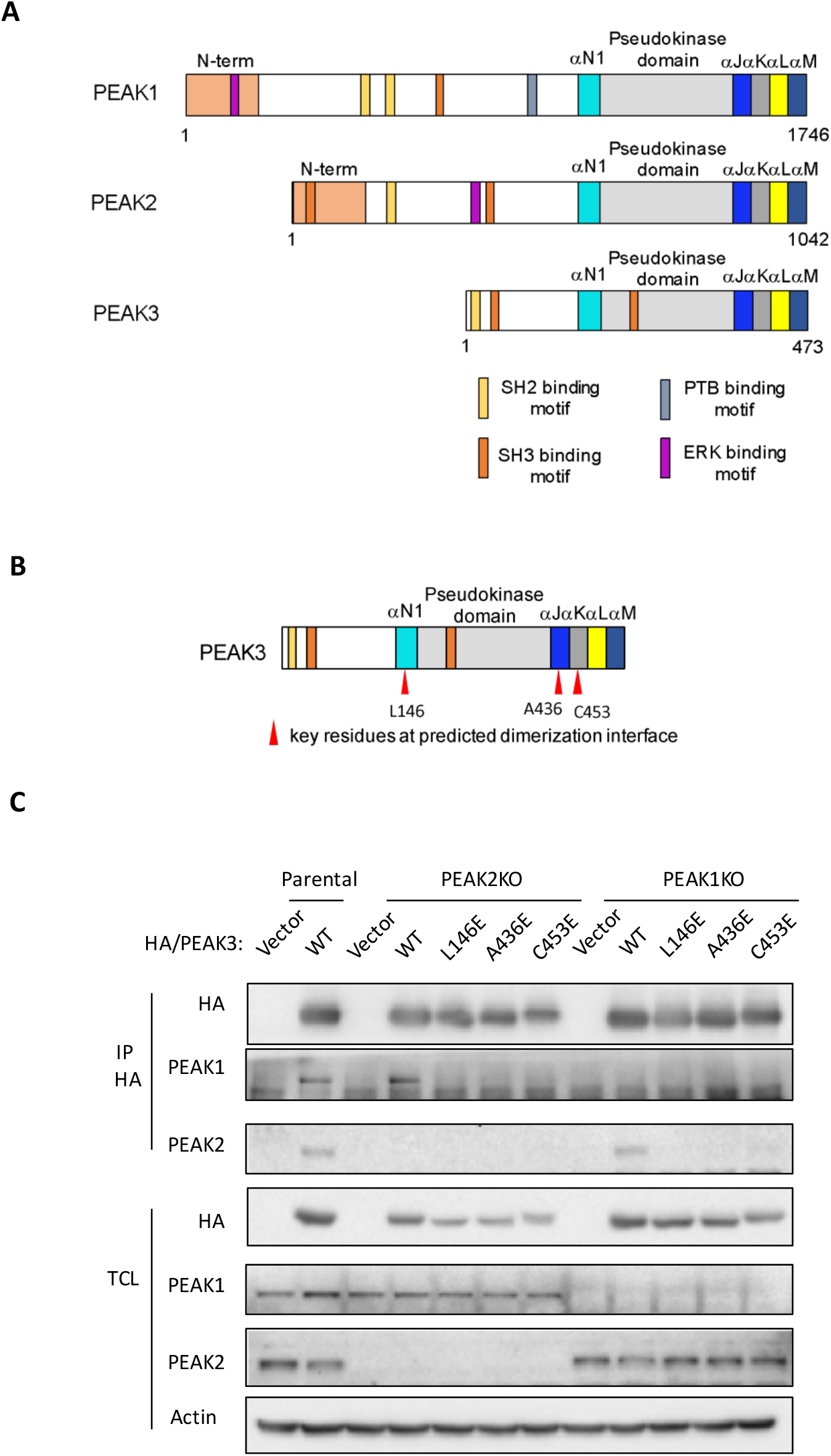
PEAK3 undergoes heterotypic association with PEAK1 and PEAK2 via the dimerization interface. A Schematic representation of PEAK1, PEAK2 and PEAK3 domain structure. B Schematic representation of key amino acid residues in predicted dimerization interface of PEAK3. C PEAK3 heterotypic association. HA-tagged WT PEAK3 or dimerization-interface mutants were overexpressed in parental, PEAK1 KO or PEAK2 KO MCF-10A cells by retroviral infection and empty vector was used as a control. Lysates were subjected to immunoprecipitation with an anti-HA antibody and Western blotted as indicated. Data are representative of three independent experiments. TCL: total cell lysate.

The presence of the conserved SHED domain in PEAK3 and demonstration that it is critical for PEAK3 homotypic association (Lopez *et al.*, 2019), led us to determine whether PEAK3 undergoes heterotypic association with PEAK1 and/or PEAK2. This was undertaken by expressing HA-tagged PEAK3 in parental MCF-10A cells. In addition, we performed similar studies in PEAK1 or PEAK2 knock-out (KO) MCF-10A cells generated by CRISPR (Fig EV1A) (Liu *et al.*, 2016; Patel *et al.*, 2017) in order to rule out ‘bridging’ of PEAK3 binding to PEAK1 by PEAK2 (and vice versa), and utilized PEAK3 SHED mutations predicted to disrupt dimerization (Fig 1B) (Lopez *et al.*, 2019) in order to confirm the underlying association mechanism. PEAK3 associated with both PEAK1 and PEAK2 in parental MCF-10A cells and interacted with PEAK1 in the absence of PEAK2, and PEAK2 in the absence of PEAK1 (Fig 1C). These data indicate that PEAK3 can form heterotypic complexes with PEAK1 or PEAK2. Furthermore, introduction of SHED mutations abolished the ability of PEAK3 to associate with PEAK1 and PEAK2 (Figs 1C and EV1B). These data indicate that PEAK3 undergoes heterotypic association with PEAK1 and PEAK2 via the conserved dimerization interface.

### Characterization of the PEAK3 scaffolding function and its regulation by EGF

The existence of SH2 and SH3 binding motifs (Fig 1A) indicates that PEAK3 may function as a scaffold, assembling protein complexes by recruiting specific binding partners. To address this hypothesis, we undertook LC-MS/MS analysis of anti-HA immunoprecipitates (IPs) prepared from MCF-10A cells expressing HA-tagged PEAK3. Proteins exhibiting significantly increased abundance in PEAK3 IPs *versus* control IPs were identified (Table EV1) and are presented in a volcano plot (Fig 2A, left panel). This confirmed PEAK3 interaction with CrkII (Lopez *et al.*, 2019), independently verified PEAK1 binding (Fig 1C) and also revealed a variety of other binding partners including the adaptors CrkL and Grb2, the ADP-ribosylation factor (ARF) GTPase-activating proteins ASAP1 and ASAP2, and non-receptor protein tyrosine kinase PYK2. In addition, since growth factor stimulation markedly impacts the assembly of PEAK1 signalling complexes (Zheng *et al.*, 2013), the effect of EGF treatment on the PEAK3 interactome was also characterized (Fig 2A, right panel and Table EV1). Interestingly, this revealed that the ability of PEAK3 to associate with ASAP1/2 and Grb2 was decreased by 5 min EGF treatment, whereas recruitment of the E3 ubiquitin ligase Cbl was increased.

**Figure 2.**
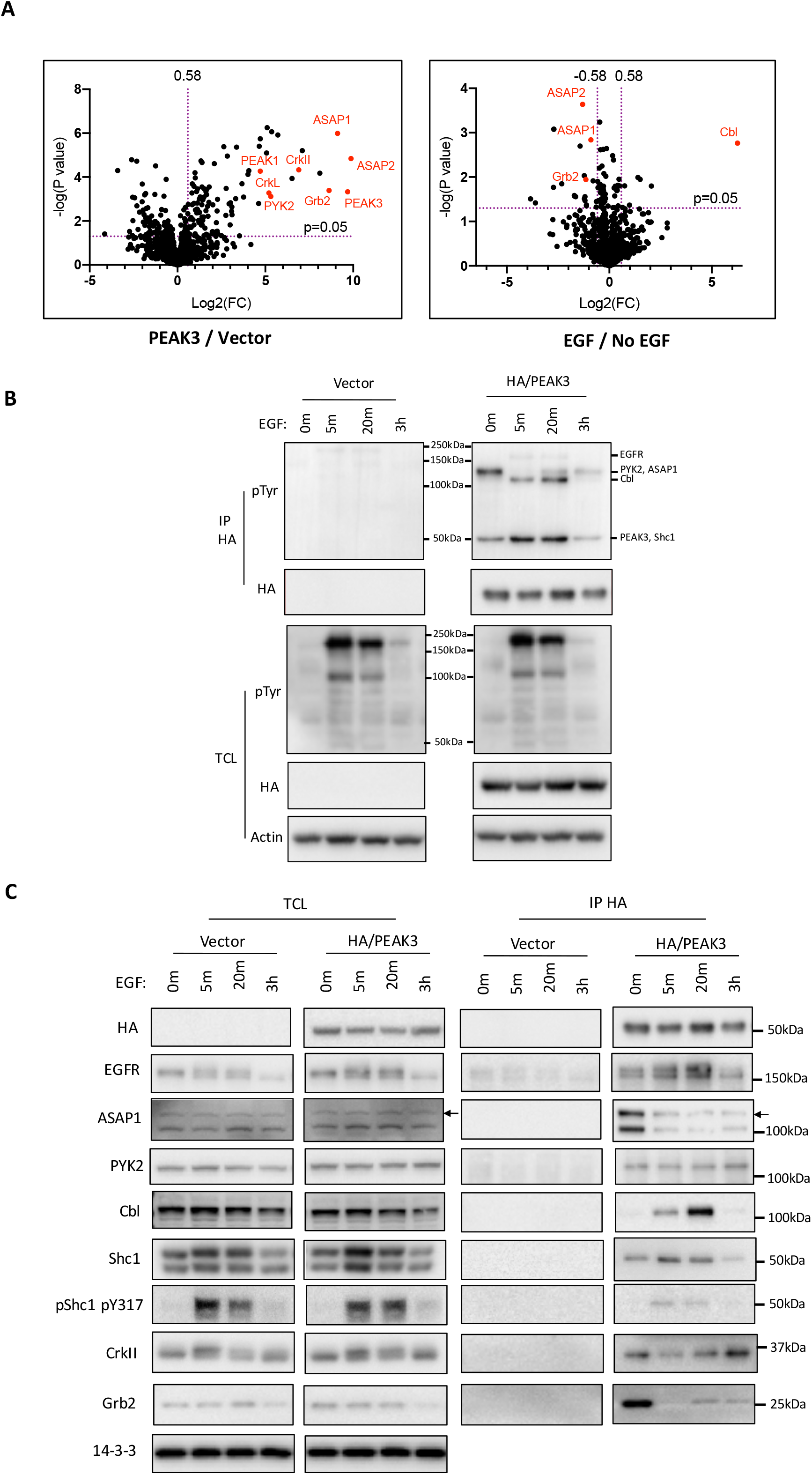
Characterization of PEAK3 binding partners. A Interactomes of PEAK3 under −/+ EGF stimulation represented as volcano plots of the MS data. MCF-10A cells stably overexpressing HA-tagged PEAK3 or empty vector were cultured in -EGF medium as described in Methods overnight and stimulated with EGF (50 ng/ml) for 5 min. Anti-HA IPs were prepared and subjected to on-beads tryptic digestion. Resulting peptides were subjected to LC-MS/MS analysis. Dashed lines indicate the p<0.05 and 1.5-fold change (Log2(FC)=0.58, up for PEAK3 versus Vector, up and down for EGF versus No EGF) thresholds. Most pronounced responding proteins are indicated in red dots and labelled. B, C Validation of MS results by Western blot. Anti-HA IPs were prepared using DKO MCF-10A cells stably overexpressing HA-tagged PEAK3 or empty vector treated with EGF for different time points, and Western blotted as indicated. Data are representative of at least three independent experiments. Predicted tyrosine phosphorylated proteins corresponding to the bands are indicated in (B).

To complement these analyses, we undertook anti-phosphotyrosine Western blotting of anti-HA IPs prepared from control or EGF-treated PEAK1/2 double KO (DKO) MCF-10A cells (Fig EV1A) expressing HA-tagged PEAK3 (Fig 2B). This detected a ~125 kDa band that was decreased by EGF treatment, and a ~50 kDa band that increased. Based on size, the defined interactome (Fig 2A) and known PEAK1 binding partners, possible candidates represent ASAP1 and either PEAK3 or Shc1, respectively. In addition, another two bands at ~175 and ~120 kDa were detected at 5 min and 20 min after EGF treatment, which may correspond to the EGFR and Cbl, respectively. Consistent with the IP/MS and anti-phosphotyrosine blotting results, IP/Westerns with specific antibodies demonstrated association of PEAK3 with Grb2, CrkII, PYK2, ASAP1 and EGFR under control conditions, and EGF treatment led to decreased association of PEAK3 with Grb2 and ASAP1 within 5 min, and increased interaction with the EGFR and Cbl, peaking after 20 min stimulation (Fig 2C). Of note, binding of CrkII was also rapidly reduced following EGF stimulation, but this returned to control levels after 20 min - 3 h EGF treatment (Fig 2C).

In order to resolve the identity of the 50 kDa tyrosine phosphorylated protein detected in PEAK3 IPs (Fig 2B), we first denatured the lysates prior to IP, so that the IP would only isolate PEAK3 and not associated proteins. Unexpectedly, this revealed that tyrosine phosphorylation of PEAK3 was downregulated by EGF treatment (Fig 3A), indicating that one or more other PEAK3 interactor(s) must contribute to the upregulated ~50 kDa pY band. A top candidate was Shc1, a well-characterized PEAK1 and Grb2 binding partner (Zheng *et al.*, 2013). Indeed, direct blotting revealed that association of both total and Y317-phosphorylated Shc1 with PEAK3 was increased by EGF treatment, reaching a maximum at 5 min and then decreasing (Fig 2C). The dynamics of the PEAK3 interactome in response to EGF treatment are summarized in Fig EV2. Collectively, these data indicate that PEAK3 functions as a scaffold and via specific downstream effectors may regulate a variety of processes and biological endpoints including cytoskeletal dynamics and cell motility, EGF downstream signalling and protein ubiquitylation and turnover.

**Figure 3.**
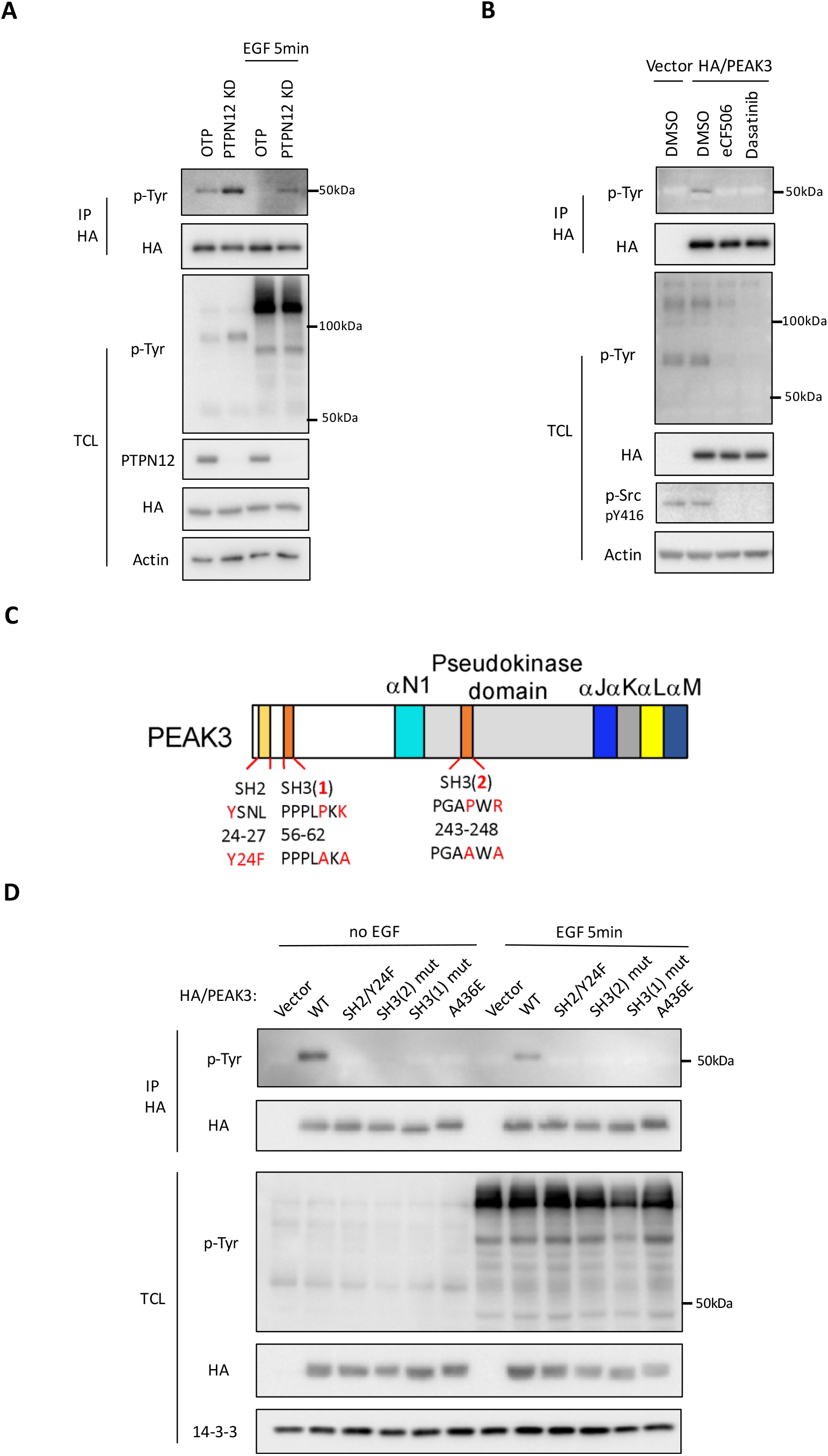
Regulation of PEAK3 tyrosine phosphorylation. A Effect of PTPN12 knockdown on PEAK3 tyrosine phosphorylation. PTPN12 siRNA pool (20 nM) was transfected into DKO MCF-10A cells stably overexpressing HA-tagged PEAK3, after 48 h of transfection the cells were starved in -EGF medium overnight and treated with EGF (50 ng/ml) for 5 min. Non-targeting siRNA pool (OTP) was used as control. B Effect of Src inhibition on PEAK3 tyrosine phosphorylation. DKO MCF-10A cells stably overexpressing HA-tagged PEAK3 were starved in -EGF medium overnight and treated with Src inhibitors eCF506 (250 nM) or Dasatinib (100 nM) for 1 h. C Schematic representation of mutated SH2 and SH3 domain binding motifs in PEAK3. D Effect of EGF treatment and mutation of SH2 and SH3 binding motifs on PEAK3 tyrosine phosphorylation. DKO MCF-10A cells stably overexpressing HA-tagged WT PEAK3 or specific mutants were starved in -EGF medium overnight and treated with EGF (50 ng/ml) for 5 min. A, B and D Anti-HA IPs were prepared from denatured cell lysates and Western blotted as indicated. Data are representative of at least three independent experiments.

### Regulation of PEAK3 tyrosine phosphorylation

Our finding that PEAK3 is rapidly dephosphorylated in response to EGF stimulation (Fig 3A and D) led us to interrogate the underlying mechanism. It was previously established that site-selective tyrosine phosphorylation of Shc1 is temporally regulated following EGF stimulation, with the protein tyrosine phosphatase (PTP) N12 dephosphorylating the Shc1/Grb2 complex prior to recruitment of PEAK1 to Shc1 to promote cell migration/invasion (Zheng *et al.*, 2013). This linkage of PTPN12 to PEAK family signalling led us to test its role in regulating PEAK3 tyrosine phosphorylation. Indeed, knockdown of PTPN12 in MCF-10A DKO cells enhanced basal phosphorylation of PEAK3 and markedly attenuated EGF-induced dephosphorylation of this scaffold, demonstrating that the latter process is PTPN12-dependent (Fig 3A). In addition, since tyrosine phosphorylation of PEAK1 is mediated by SFKs (Croucher *et al.*, 2013; Wang *et al.*, 2010), we characterized the role of these kinases. Treatment of MCF-10A DKO cells with either the selective SFK inhibitor eCF506 (Fraser *et al*, 2016) or the SFK/Abl inhibitor dasatinib blocked PEAK3 tyrosine phosphorylation, indicating that this modification is dependent on SFK activity (Fig 3B).

In order to determine the role of dimerization and specific SH2 and SH3 binding motifs in PEAK3 tyrosine phosphorylation, HA-tagged PEAK3 proteins with mutations in the dimerization interface and specific SH2 and SH3 binding motifs (Figs 1B and 3C) were expressed in MCF-10A DKO cells. IP/Western blotting analysis using denatured lysates demonstrated that dimerization is required for efficient PEAK3 tyrosine phosphorylation (Fig 3D). Mutation of Y24 also abolished PEAK3 tyrosine phosphorylation (Fig 3D), indicating that this is the main phosphorylated site and Y_24_SNL represents a potential SH2 binding motif that conforms to a Grb2 SH2 binding consensus (pYXN) (Miller *et al*, 2008). In addition, mutation of either SH3 binding motif also prevented PEAK3 tyrosine phosphorylation (Fig 3D), highlighting a potential role in recruiting or regulating the upstream tyrosine kinase.

### Structural requirements for effector recruitment by PEAK3

Next, we turned to the PEAK3 interactome, and having demonstrated that PEAK3 dimerization is required for PEAK3 tyrosine phosphorylation (Fig 3D), determined how dimerization potential affects recruitment of specific binding partners. Flag-tagged wildtype (WT) PEAK3 was co-expressed in HEK293 cells with HA-tagged versions of WT PEAK3 or dimerization-defective mutants (Fig 4A). IP/Westerns confirmed the inability of these mutants to undergo homotypic association, and also revealed that recruitment of Grb2, CrkII, ASAP1 and PYK2 to PEAK3 was PEAK3 dimerization-dependent (Fig 4A). However, association of EGFR with these mutants was similar to that for wildtype PEAK3 (Fig 4A). These findings build significantly on the previous report from the Jura laboratory reporting dimerization-dependent CrkII association (Lopez *et al.*, 2019), highlighting a much broader impact of dimerization on assembly of the PEAK3 interactome, and also demonstrate that binding of PEAK3 monomer to certain interactors can occur.

**Figure 4.**
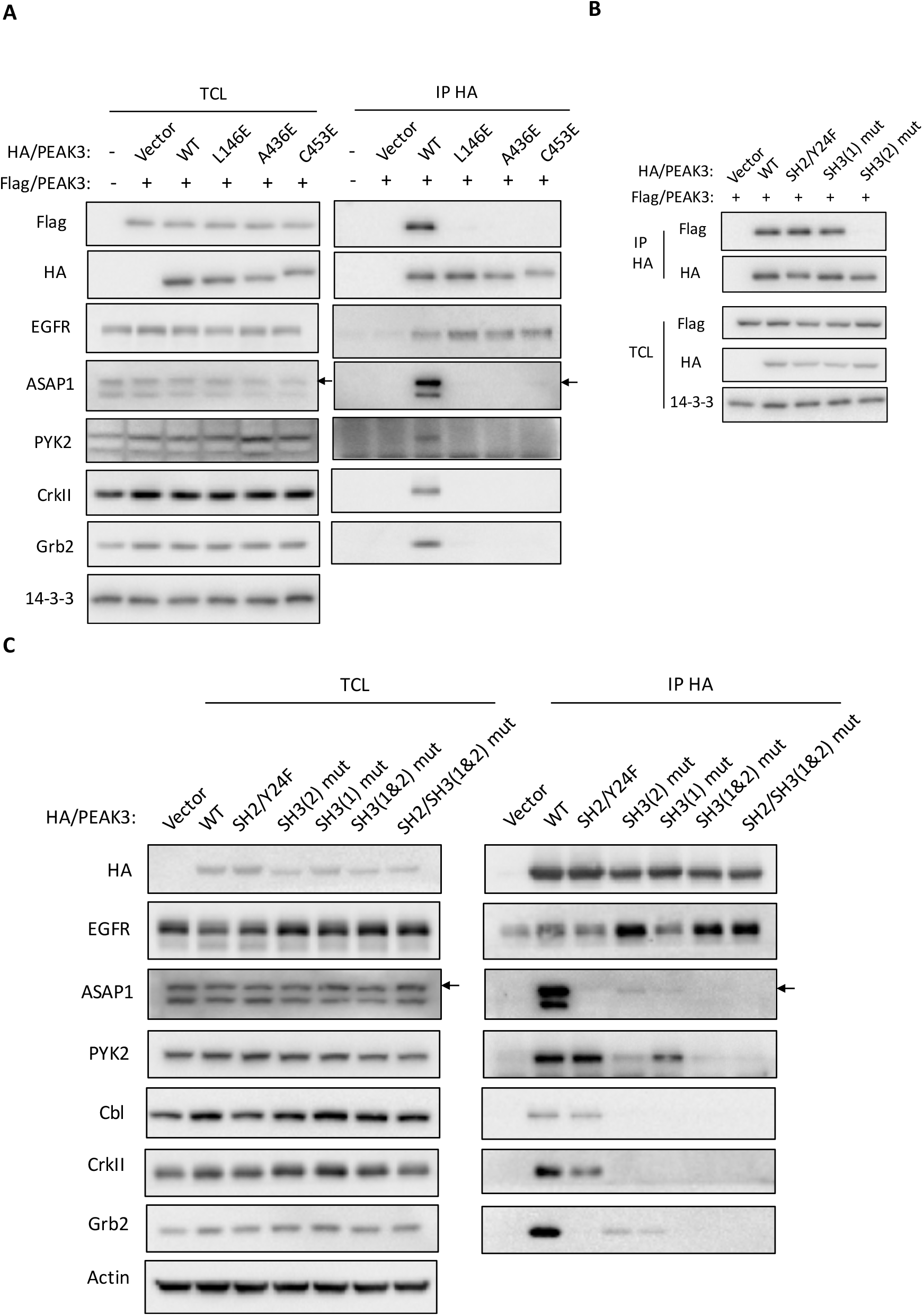
PEAK3 dimerization and interaction with binding partners. A Effect of dimerization on PEAK3 association with specific binding partners. HEK293 cells were co-transfected with plasmids encoding Flag-tagged WT PEAK3 in combination with plasmids encoding HA-tagged WT PEAK3 or dimerization-deficient mutants. B Effect of mutated SH2 and SH3 domain binding motifs on PEAK3 dimerization. HEK293 cells were co-transfected with plasmids encoding Flag-tagged WT PEAK3 in combination with plasmids encoding HA-tagged WT or specific mutant versions of PEAK3. C Role of PEAK3 SH2 and SH3 domain binding motifs in PEAK3 association with specific binding partners. HA-tagged WT PEAK3 or specific mutants were overexpressed in DKO MCF-10A cells by retroviral infection. A, B and C Empty vector was used as a control. Lysates were subjected to immunoprecipitation with an anti-HA antibody and Western blotted as indicated. Data are representative of three independent experiments.

The roles of specific SH2/SH3 binding motifs were then characterized. However, in order to correctly interpret these data, it was first necessary to determine whether mutation of these motifs impacted PEAK3 dimerization. Importantly, while mutation of the SH2 binding motif at Y24 and the N-terminal motif SH3(1) had no effect, mutation of SH3(2), which is located in the N-lobe of the PsK domain (Fig EV3), from PGAPWR to PGAAWA significantly disrupted PEAK3 dimerization (Figs 4B and EV4A). Interestingly, the SH3(2) motif is not conserved within the PEAK family and corresponds to the loop between the β4 and β5 strands that in the PEAK1 and PEAK2 crystal structures adopts an α-turn. This region also marks the start of a unique insertion loop in PEAK1 and PEAK2 that is absent in PEAK3. Even though the β4-β5 loop is located away from the dimerization domain, the SH3(2) mutation may affect the conformation in this region and thereby impact on the orientation of the N-lobe with respect to the C-lobe, the positioning of the αN1 helix and hence dimerization (Fig EV3).

With regard to recruitment of binding partners, mutation of the PEAK3 SH2 binding motif at Y24 abolished PEAK3 association with Grb2 and ASAP1 and decreased association with CrkII to approximately 40% of control. However, it had no effect on Cbl, PYK2 and EGFR binding (Figs 4C and EV4B). Disruption of SH3(1) eliminated CrkII and Cbl association, markedly reduced Grb2 and ASAP1 binding, decreased PYK2 recruitment to approximately 50%, and did not affect EGFR association. In the case of SH3(2), mutation led to undetectable, or significantly reduced, binding of Grb2, CrkII, ASAP1 PYK2 and Cbl, while association with the EGFR was enhanced (Figs 4C and EV4B). However, additional effects of the SH3 binding site mutations must be considered when interpreting these data. First, integrity of SH3(1) is required for Y24 phosphorylation (Fig 3D). Consequently, mutation of SH3(1) will indirectly reduce Y24 phosphorylation and thereby prevent SH2 domain-mediated interactions at this site, as well as direct SH3 domain-mediated binding to SH3(1). Second, since mutation of SH3(2) negatively impacts both PEAK3 dimerization and tyrosine phosphorylation, this explains the marked reduction in binding partner interactions, although the increased association with EGFR may reflect an altered conformation of the PsK domain. Integrating these data with the dynamics of Y24 phosphorylation and Grb2 and ASAP1 binding in response to EGF (Figs 2C and EV2), and considering the modular structure of Grb2 and ASAP1 (Lowenstein *et al*, 1992; Tanna *et al*, 2019), it appears likely that Grb2 binds Y24 via its SH2 domain and bridges PEAK3 to ASAP1 (Fig EV4C). The requirement for SH3(1) in ASAP1 binding may reflect its role in indirectly mediating Y24 phosphorylation, binding of SH3(1) to the ASAP1 SH3 domain, or bridging by another adaptor. With regard to the latter mechanism, although the dynamics of ASAP1 recruitment to PEAK3 are closer to those of Grb2 than CrkII (Fig. EV2), it remains possible that Grb2 and CrkII function co-operatively to recruit ASAP1 (Fig EV4D). The mechanisms of recruitment for PYK2, Cbl and EGFR require further characterization, since they do not follow the same kinetics as Grb2 or CrkII binding.

### Y24/Grb2/ASAP1 represents a key PEAK3 signalling axis

Our findings that EGF-induced dissociation of Grb2 and ASAP1 from PEAK3 follow similar kinetics and occurs concurrent with PEAK3 dephosphorylation, and that binding of both proteins to PEAK3 is lost upon Y24 mutation (Figs 2–4), highlight the importance of Y24 phosphorylation to PEAK3 signalling. In order to interrogate this potential signalling axis in more detail, we undertook Isothermal Titration Calorimetry (ITC) experiments to measure the binding of recombinant full-length Grb2 or CrkII to synthetic 7-mer peptides, either tyrosine phosphorylated or non-phosphorylated, encompassing the Y24 SH2 binding motif in PEAK3. This confirmed direct and specific binding of both adaptors to the phosphopeptide with 1:1 stoichiometry, with Grb2 exhibiting a 3-fold higher affinity for this site (measured dissociation constants, *K*_D_, of the phosphopeptide were 2.6 μM and 7.8 μM for Grb2 and CrkII respectively) (Fig 5A-C), consistent with the greater role for this site in Grb2 recruitment (Fig 4C). To determine the requirement for Grb2 in assembly of the PEAK3 interactome, we knocked down Grb2 then undertook Western blot analysis of PEAK3 IPs. Strikingly, this resulted in loss of ASAP1 binding, while association with PYK2 was unaffected (Fig 5D). However, it also led to undetectable PEAK3 tyrosine phosphorylation and reduced CrkII recruitment (Fig 5D), the latter consistent with the effect of Y24 mutation (Fig 4C). These data indicate that ASAP1 recruitment to PEAK3 is indeed Grb2-dependent, but because PEAK3 tyrosine phosphorylation is also dependent on the presence of Grb2, this may reflect Grb2-dependent Y24 phosphorylation, bridging by Grb2, or both. Nonetheless, they reveal a key PEAK3 signalling axis that can be further characterized in terms of its functional role.

**Figure 5.**
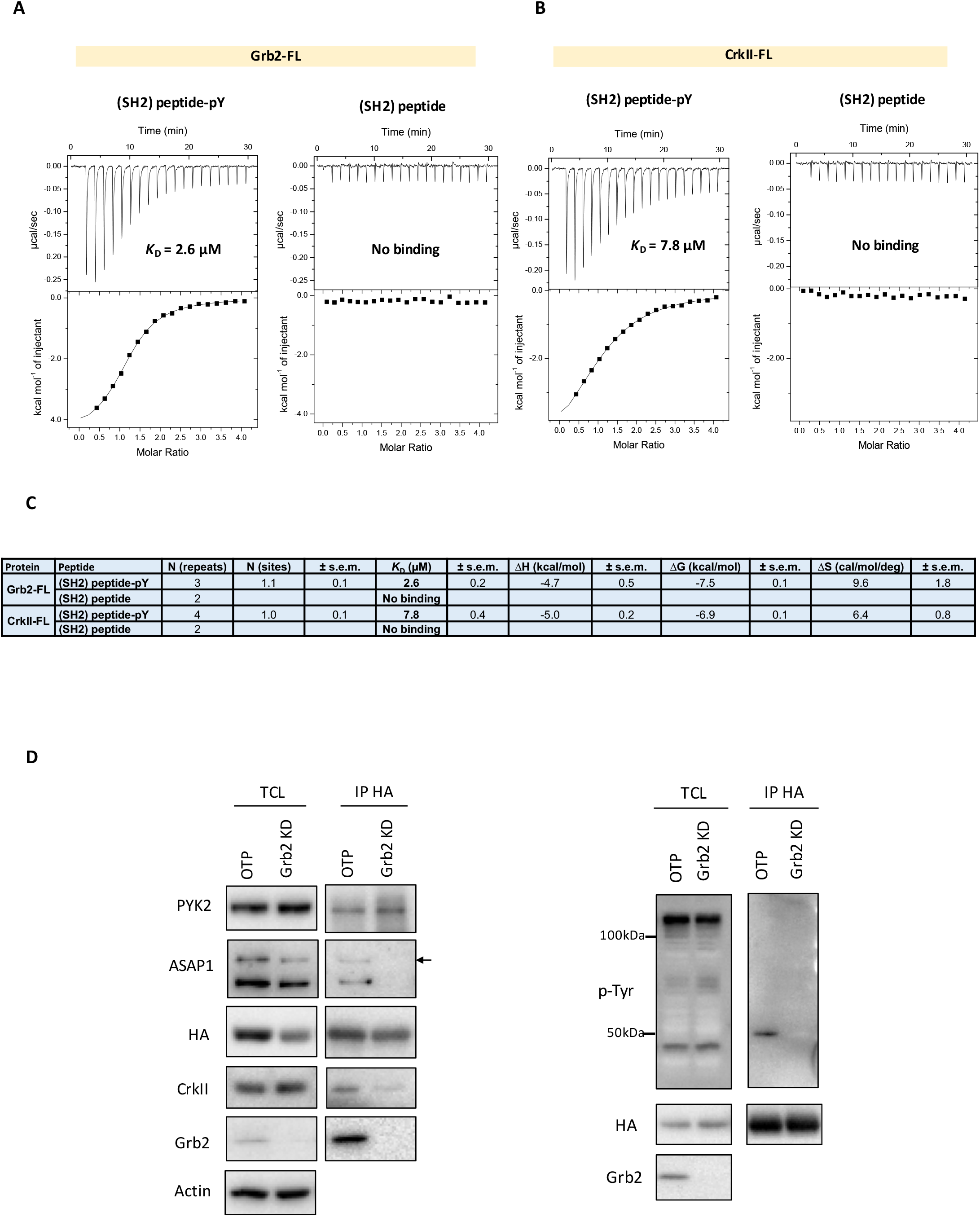
PEAK3 binding with Grb2 and CrkII. A ITC data for binding of PEAK3 phosphorylated peptide (T(pY)SNLGQ) and non-phosphorylated peptide (TYSNLGQ) with full-length Grb2 (Grb2-FL). The equilibrium dissociation constant (K_D_) for the interaction of Grb2 with the PEAK3 peptide-pY was 2.6 ± 0.2 μM from three replicate experiments. B ITC data for the binding of PEAK3 phosphorylated peptide (T(pY)SNLGQ) and non-phosphorylated peptide (TYSNLGQ) with full-length CrkII (CrkII-FL). The K_D_ of the interaction of CrkII with the PEAK3 peptide-pY was 7.8 ± 0.4 μM from four replicate experiments. C Data of A and B were fitted to a single binding site model to obtain the stoichiometry (N), the dissociation constant (K_D_) and the enthalpy of binding (ΔH). ΔG is Gibbs free energy change on binding, ΔS is the entropy change on binding. The reported values are the mean ± s.e.m. from independent measurements as indicated. D Effect of Grb2 knockdown on PEAK3 association with specific binding partners and tyrosine phosphorylation. Grb2 siRNA pool (20 nM) was transfected into DKO MCF-10A cells stably overexpressing HA-tagged PEAK3. Non-targeting siRNA pool (OTP) was used as control. Anti-HA IPs were prepared and Western blotted as indicated. In the right-hand panel, IPs were prepared from denatured lysates. Data are representative of at least three independent experiments.

### PEAK3 promotes cell elongation and motility

While PEAK1 and PEAK2 positively regulate cell elongation and migration (Bristow *et al*, 2013; Croucher *et al.*, 2013; Safari *et al.*, 2011; Tactacan *et al.*, 2015), PEAK3 blocks CrkII-dependent membrane ruffling, suggesting a possible antagonistic role in PEAK family signalling (Lopez *et al.*, 2019). To characterize the functional role of PEAK3, WT PEAK3 or specific PEAK3 mutants were stably expressed in MDA-MB-231 breast cancer cells (Fig 6A). Expression of WT PEAK3 led to clear transition to a more elongated morphology (Fig 6B and C), but this effect was lost with introduction of the dimerization disrupting mutations A436E and SH3(2) (Fig 6C).

**Figure 6.**
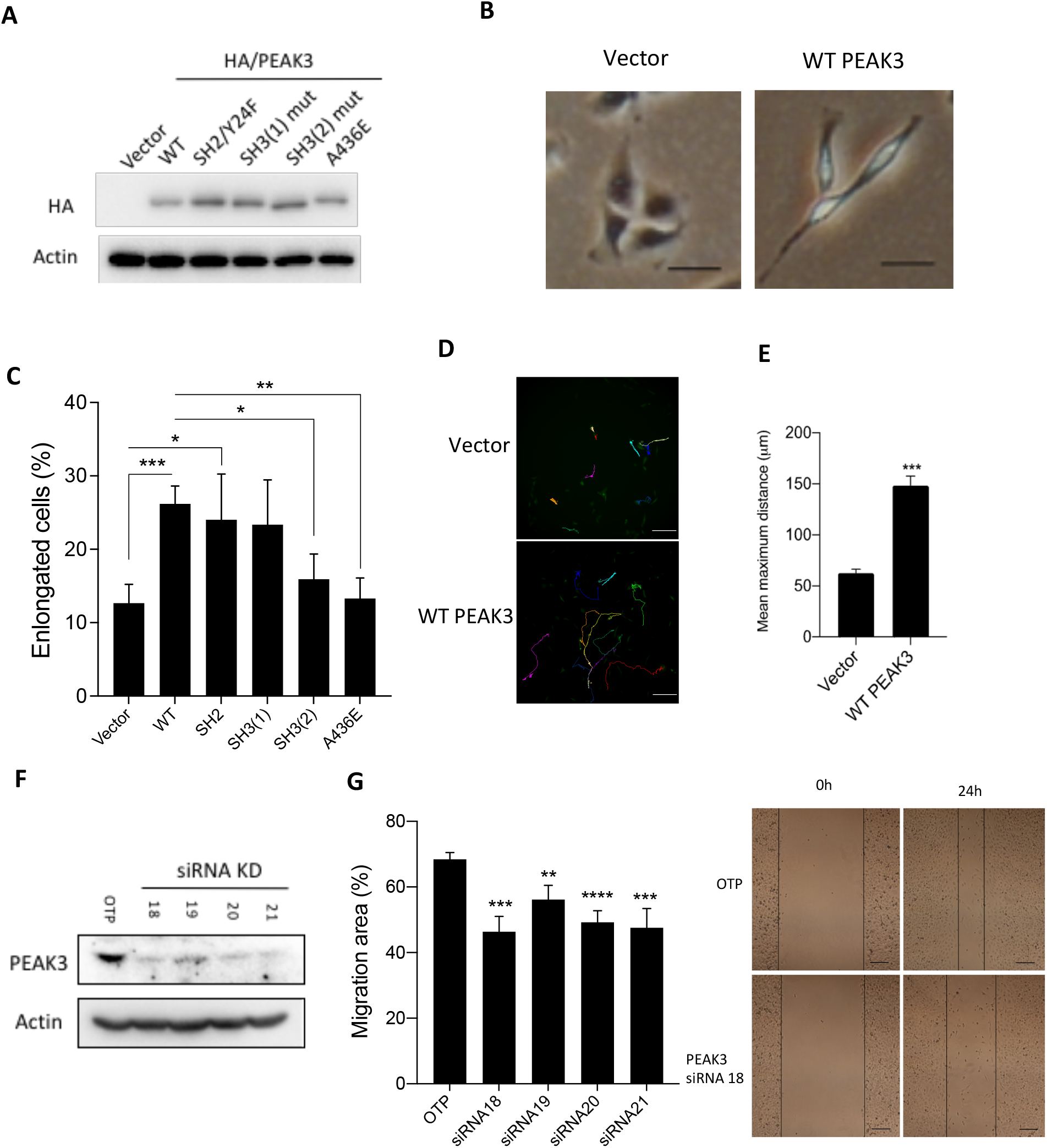
Effect of PEAK3 on elongation and motility in MDA-MB-231 cells. A Expression levels of HA/PEAK3 and mutants in MDA-MB-231 cells. Lysates prepared from cells stably expressing HA-tagged WT PEAK3 and mutants were Western blotted as indicated. B, C Effect of WT PEAK3 and different mutants on elongation in MDA-MB-231 cells. MDA-MB-231 cells in (A) were used for morphology analysis and quantification of the percentage of elongated cells. Representative images of WT PEAK3-expressing cells are shown in (B). Scale bar, 50 μm. Data in (C) represent the mean ± s.e.m. of three independent experiments. At least 1000 cells were analyzed in each experiment. D, E Effect of WT PEAK3 on random cell motility in MDA-MB-231 cells. HA-tagged WT PEAK3 was stably overexpressed in MDA-MB-231 cells and random cell motility was determined by live cell tracking. Representative images are shown in (D). The histogram in (E) indicates the average maximum displacement distance from the origin. Scale bar, 200 μm. Data represent the mean ± s.e.m. of three independent experiments. ~60 cells were analyzed in each experiment. F, G Knockdown of endogenous PEAK3 inhibits MDA-MB-231 cell migration. PEAK3 individual siRNAs 18-21 (20 nM) were transfected into MDA-MB-231 cells. Cell lysates were Western blotted (F) and a wound healing assay was performed (G). OTP was a non-targeting control. The histogram in (G) provides the quantitative analysis of the relative wound area for 4 different PEAK3 siRNAs. *, p<0.05, **, p<0.01, ***, p<0.001, ****, p<0.0001, by Student’s t-test. The representative images in (G) were taken immediately after scratches had been made and after 24 h incubation. The black lines indicate the wound area. Scale bar, 200 μm.

The role of PEAK3 in regulating cell migration was also characterized. First, we established that overexpression of PEAK3 in MDA-MB-231 cells leads to enhanced random cell motility, as determined by live cell tracking (Fig 6D and E). Next, we characterized the role of endogenous PEAK3. In order to undertake this study, we first generated a custom polyclonal anti-PEAK3 antibody. This detected ectopically-expressed PEAK3 (Fig EV5A) and an approximately 50 kDa band in MDA-MB-231 cell lysates that was knocked down by each of 4 independent siRNAs directed against PEAK3 (Fig 6F) and ablated by CRISPR-mediated knockout (Fig EV5B). Endogenous PEAK3 was also detected in RH30 sarcoma and A172 glioblastoma cells (Fig EV5B and C). Having established this assay for PEAK3 protein expression, we characterized the effect of PEAK3 knockdown. Importantly, use of each of the PEAK3-directed siRNAs significantly reduced MDA-MB-231 migration in a scratch assay (Fig 6G), confirming the pro-migratory role of endogenous PEAK3.

### PEAK3 promotes growth and invasion of MCF-10A acini in three-dimensional culture

Growth of MCF-10A breast epithelial cells in 3D Matrigel culture represents a powerful model system for characterizing the effect of specific signalling proteins on a variety of biological endpoints, including proliferation, apoptosis, polarity and invasion (Debnath *et al*, 2003), and was used to identify PEAK1 as a breast cancer oncogene (Croucher *et al.*, 2013). Therefore, we exploited this model to provide further insights into PEAK3 function. In addition, we utilized DKO MCF-10A cells, in order to interrogate PEAK3 in the absence of PEAK1 and PEAK2. Overexpression of PEAK3 led to a significant increase in acinar size (Fig 7A-C) at Day 5 and Ki67-positive acini at Day 12 (Fig 7D and E), highlighting for the first time a pro-proliferative role for PEAK3 and indicating that expression of this scaffold can overcome the proliferative suppression that normally occurs in late-stage MCF-10A cultures (Debnath *et al.*, 2003). This effect of PEAK3 on acinar growth was reduced for all of the mutants analysed (Fig 7C).

**Figure 7.**
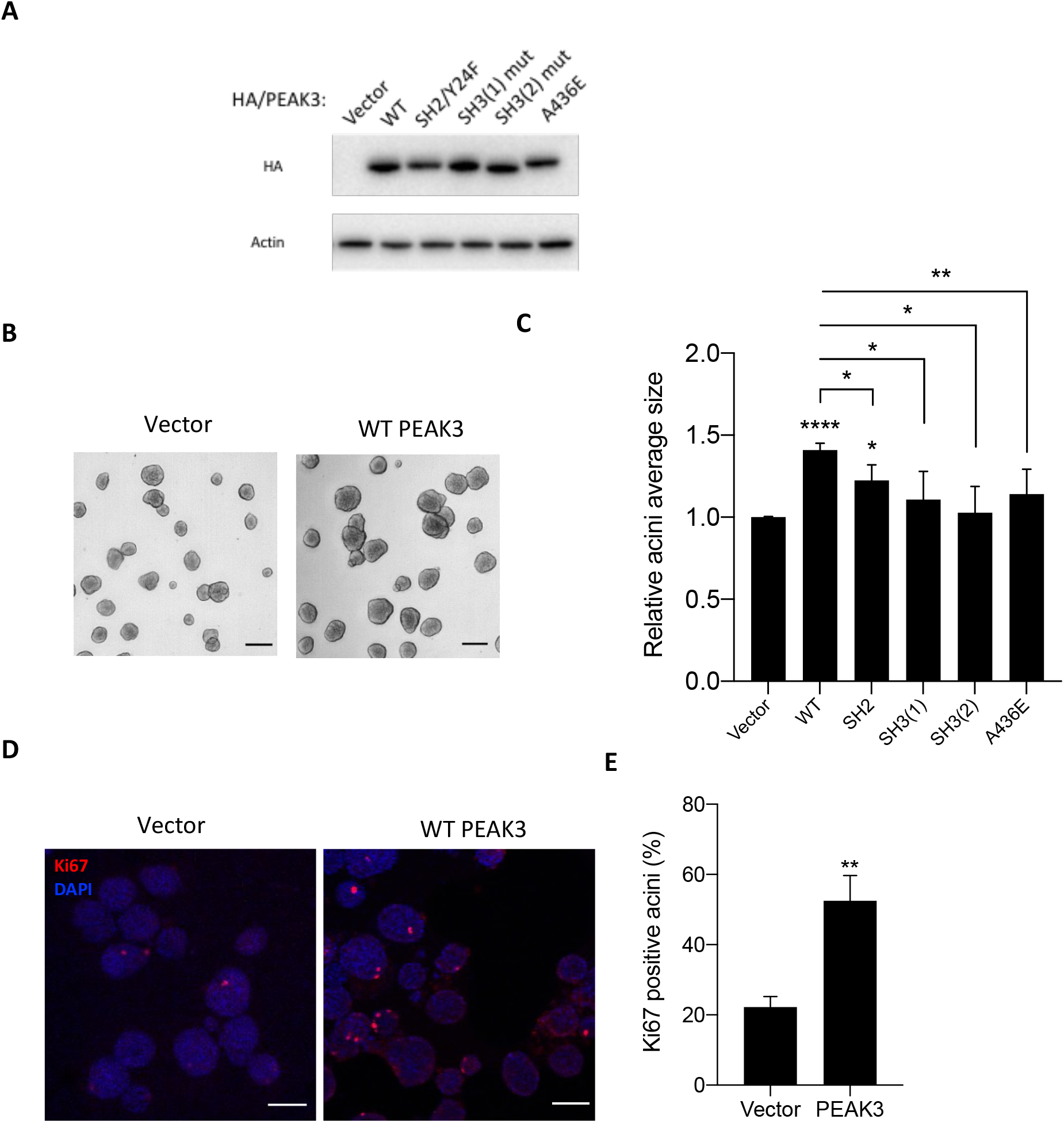
Effect of PEAK3 on MCF-10A acini growth. A Expression levels of HA/PEAK3 and mutants in DKO MCF-10A cells. Lysates prepared from stably expressing cells were Western blotted. B, C Effect of WT PEAK3 and specific mutants on MCF-10A acini growth. DKO MCF-10A cells in (A) were used for 3D culture. Micrographs, acini grown on Matrigel for 5 days. Scale bar, 100 μm. Histogram, quantification of the relative average size of acini expressing PEAK3 WT and the different mutants. D, E Fluorescence microscopy of acini grown on Matrigel for 12 days and stained with Ki67 antibody (red) and nuclei visualized with DAPI (blue), subsequently analysed by confocal microscopy. Histogram, quantification of Ki67 positive acini. Data represent the mean ± s.e.m. of 3 independent experiments. ≥50 acini were analyzed in each experiment. *, p<0.05, **, p<0.01, ****, p<0.0001, by Student’s t-test.

Interestingly, an additional phenotype observed in late stage PEAK3-overexpressing cultures was cell invasion, characterized by a failure in basal membrane integrity leading to cells invading the surrounding matrix (Fig 8A-B). Strikingly, this phenotype was dependent on not only PEAK3 dimerization, but also the two SH3 binding sites and Y24 (Fig 8C). Since the latter is critical for not only Grb2 binding but also recruitment of ASAP1, this highlights an important role for the Y24/Grb2/ASAP1 signalling axis in PEAK3-mediated cell invasion.

**Figure 8.**
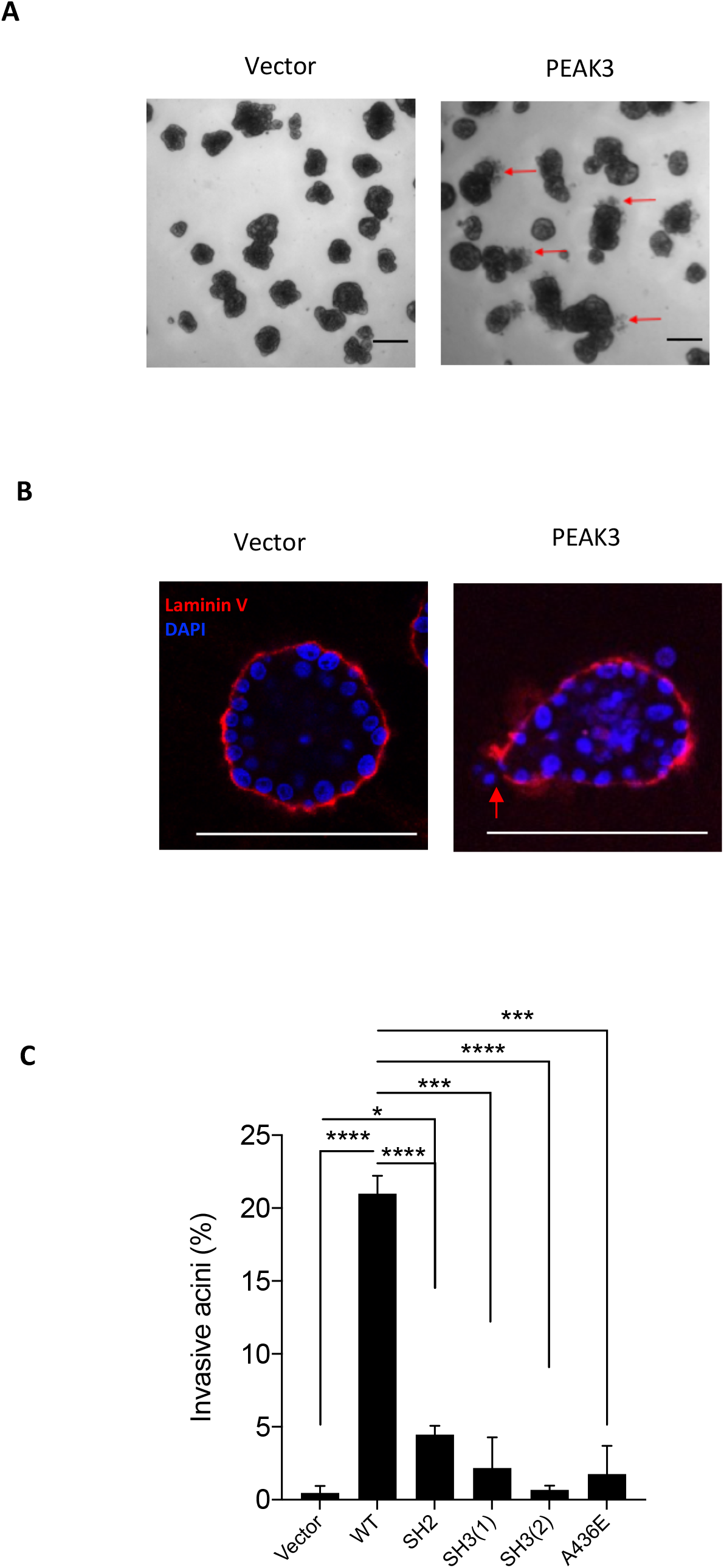
Effect of PEAK3 on MCF-10A acini invasion. A Effect of WT PEAK3 on MCF-10A acini invasion. Micrographs of vector control and PEAK3 overexpressing DKO MCF-10A acini grown on Matrigel for 12 days. Red arrowheads indicate invasive cells. Scale bar, 100 μm. B Fluorescence microscopy indicate acini with invasive phenotype, stained with antibody against laminin V (red) and nuclei visualized with DAPI (blue), and subsequently analysed by confocal microscopy. Red arrowheads indicate invasive cells. Scale bar, 100 μm. C Effect of WT PEAK3 and specific mutants on MCF-10A acini invasion. DKO MCF-10A cells overexpressing WT PEAK3 and specific mutants were maintained in 3D culture for 12 days. The histogram indicates the quantification of the number of acini with invasive phenotype per 100 acini. Data represent the mean ± s.e.m. of three independent experiments. *, p<0.05, ***, p<0.001, ****, p<0.0001, by Student’s t-test.

## Discussion

The pseudokinases PEAK1 and 2 have emerged as key regulators of cell migration and signalling in both normal and cancer cells (Patel *et al.*, 2020). Based on a shorter N-terminal extension and ability to block CrkII-enhanced membrane ruffling and cell morphology change, it was recently proposed the new family member PEAK3 might represent a negative regulator that antagonizes PEAK1/2 function (Lopez *et al.*, 2019). However, in this study, through detailed mechanistic and functional characterization, including knockdown of the endogenous protein, we determine that PEAK3 instead represents a positive-acting scaffold that significantly enhances the signalling repertoire and output of the PEAK family and promotes cell proliferation, migration and invasion. In addition, the characterization of PEAK3 reveals complex mechanisms underpinning effector recruitment with significant implications for understanding signalling by the PEAK pseudokinases as well as scaffolds and adaptors more generally.

An important finding was that despite its shorter N-terminal extension, PEAK3 recruits several proteins not bound by PEAK1/2, including Cbl, PYK2 and ASAP1. Since we demonstrate that PEAK3 can undergo homo-as well as heterotypic association, these data indicate that PEAK3 has important independent signalling functions and can also expand the signalling potential of the PEAK family by associating with PEAK1/2 and assembling signalling complexes with contrasting outputs. This expansion of signalling repertoire via dimerization is also observed in the erbB family of receptor tyrosine kinases (Citri & Yarden, 2006), although in this case only one family member is a pseudokinase (erbB3), in contrast to the PEAK family where this is a common feature. With regard to binding of the E3 ubiquitin ligase Cbl by PEAK3, this might represent a mechanism for promoting ubiquitin-mediated degradation of not only PEAK3 itself but also PEAK1/2 bound as partners in a heterodimer. In addition, since PEAK3 undergoes EGF-induced association with the EGFR, this might also represent a mechanism for promoting EGFR ubiquitylation. Furthermore, given the known roles of the PEAK family in regulating cytoskeletal organization and cell migration/invasion, the identification of PYK2 and ASAP1 as PEAK3 binding partners identifies important downstream effectors that may mediate these effects. In this regard, binding of ASAP1 provides a potential mechanism to promote cell invasion, since ASAP1 enhances podosome and invadopodia formation in transformed cells (Bharti *et al*, 2007). Indeed, we demonstrate that recruitment of ASAP1 by PEAK3 is required for the latter to promote a novel invasive phenotype in MCF-10A cells in 3D culture.

Consistent with a previous study (Lopez *et al.*, 2019), we determined that dimerization is required for CrkII binding by PEAK3. However, a striking finding was that this requirement extended more generally to the scaffolding function of PEAK3, with recruitment of Grb2, ASAP1 and PYK2 to PEAK3 also lost upon mutation of the PEAK3 dimerization interface. A detailed understanding of this will require structural studies, but it seems likely that a number of mechanisms underpin this finding. First, the dependency of PEAK3 Y24 phosphorylation on dimerization, since the Y24 SH2 binding site is required for optimal CrkII/Grb2 binding. Here a similar mechanism to that regulating PEAK2 tyrosine phosphorylation may apply, where dimerization activates an associated tyrosine kinase, which in the case of PEAK2 is Csk (Lecointre *et al.*, 2018; Patel *et al.*, 2017; Senda *et al.*, 2016). Based on mutational studies, binding of this kinase to PEAK3 is directly or indirectly mediated via SH3(1). Second, dimerization may lead to generation of binding interfaces not present in the individual monomers. Third, the ability of the bridging adaptors to dimerize, which has been demonstrated for Grb2 (Benfield *et al*, 2007), and is likely to occur with CrkII, based on studies on the closely-related CrkL (Harkiolaki *et al*, 2006). In this binding mode, significant co-operativity is likely to be observed for binding of the adaptors to the PEAK3 dimer versus the monomer. In addition, it should be noted that particular downstream effectors can also dimerize, including Cbl (Shivanna *et al*, 2015) and ASAP1 (Chen *et al*, 2020).

PEAK1 qualitatively regulates tyrosine kinase signal output, mediating a ‘switch’ in Shc1 signalling from mitogenic/survival signalling at early time points following EGF stimulation to control of cell morphology/migration at later times (Zheng *et al.*, 2013). This ‘switch’ also involves PTPN12, which dephosphorylates the early phase Shc1/Grb2 complex prior to recruitment of PEAK1 to Shc1. In addition, PEAK1 exerts quantitative effects on signalling, promoting aberrant proliferation, migration and invasion when overexpressed (Croucher *et al.*, 2013) and exhibiting upregulation in several cancer types (Agajanian *et al.*, 2015; Croucher *et al.*, 2013; Ding *et al.*, 2018; Huang *et al.*, 2018; Kelber *et al.*, 2012). Importantly, our study reveals interesting similarities between PEAK1 and PEAK3 that extend beyond dimerization and adaptor protein recruitment. Upon EGF treatment, PEAK3 was rapidly dephosphorylated in a PTPN12-dependent manner, highlighting an important role for this PTP in regulating assembly of PEAK family signalling complexes. Since PEAK3 tyrosine phosphorylation regulates Grb2/ASAP1 recruitment, this is consistent with establishing proliferative/survival signals rather than pro-migratory ones during the early phase of EGF receptor signalling (Zheng *et al.*, 2013). In addition, PEAK3 can promote aberrant signalling when overexpressed, leading to failure in the basement membrane integrity of MCF-10A acini and cellular invasion into the surrounding matrix. Given this finding, it will be important to characterize PEAK3 expression in specific cancers and determine whether its downstream pathways represent therapeutic targets.

## Materials and Methods

### Plasmids

Codon-optimized cDNAs encoding N-terminal HA or Flag-tagged WT PEAK3 and HA-tagged PEAK3 mutants were synthesized by Genscript and cloned into the EcoRI restriction sites of the pMIG-GFP Express vector.

### Antibodies and reagents

The following antibodies were used in the study: The PEAK2 antibody was custom-made (Tactacan *et al.*, 2015), as was the PEAK3 antibody (Cambridge, Project Code: CRB240, antigen: SSPEPPTEPPEPDNPTW), PEAK1 (Santa Cruz Biotechnology, cat. sc-100403); HA (Cell Signalling, cat. 3724), Flag (Sigma, cat. F1804), p-Tyr (Cell Signalling, cat. 8954S), PYK2 (Cell Signalling, cat. 3480S), Cbl (Santa Cruz Biotechnology, cat. sc-170), EGFR (Cell Signalling, cat. 4267S), CrkII (Santa Cruz Biotechnology, cat. sc-289), Shc (BD Biosciences, cat. 610879), p-Shc (Cell Signalling, cat. 2431S), ASAP1 (Santa Cruz Biotechnology, cat. sc-11539), Ki67 (Cell Signalling, cat. 9027), 14-3-3 (Santa Cruz Biotechnology, cat. sc-1657), PTPN12 (Abcam, cat. ab76942), p-Src Y416 (Cell Signalling, cat. 2101S), β-actin (Santa Cruz Biotechnology, cat. sc-69879), Laminin V (Chemicon, cat. MAB19562). AlexaFluor-conjugated (Invitrogen) and HRP-linked (Bio-Rad) secondary antibodies against rabbit and mouse IgG were used in Immunofluorescence staining and Western blotting, respectively. eCF506 was obtained from Professor Margaret Frame (Edinburgh Cancer Research Centre) (Fraser *et al.*, 2016). Dasatinib was purchased from Bristol Myers Squibb (BMS-354825-03). Two synthetic 7 mer peptide variants, either phosphorylated or non-phosphorylated at the key tyrosine residue, were purchased from Mimotopes (pY: Ac-T(pY)SNLGQ-NH2; Y: Ac-TYSNLGQ-NH2; N-terminally acetylated, C-terminally amidated; purity ≥ 95%).

### Tissue culture and generation of stable cell lines

MDA-MB-231 cells were obtained from EG&G Mason Research Institute, Worcester, MA and maintained in RPMI-1640 medium (Gibco, cat. 31800089) supplemented with 10% FBS (Moregate, cat. MORFBSFAU, Batch: 50301112), 10◻μg/mL Actrapid penfill insulin (Clifford Hallam Healthcare), and 20◻mM HEPES (Life Technologies, cat. 15630080). The MCF-10A cell line stably expressing the murine ecotropic receptor (MCF-10A EcoR) was a kind gift from Drs. Danielle Lynch and Joan Brugge, Harvard Medical School. MCF-10A EcoR cells were maintained as previously described (Brummer *et al*, 2006). RH30 cells were maintained in RPMI-1640 medium supplemented with 10% FBS (Moregate). HEK293, PlatE, A172 and SK-N-BE2 cells were maintained in DMEM (Gibco, cat.1200046) supplemented with 10% FBS (Moregate). PEAK1 and PEAK2 knock-out (KO) MCF-10A cells were generated by CRISPR as previously described (Liu *et al.*, 2016) and double knock-out (DKO) cells were generated by knocking out PEAK1 in PEAK2 KO cells. PEAK3 knock-out (KO) MDA-MB-231 and RH30 cells were generated by CRISPR. The single guide RNAs targeting PEAK3 were designed using GPP sgRNA Designer (Doench *et al*, 2016) and the sequences were gRNA1: CGCCTCGGAGACCCTTTCTG, gRNA2: GCCCACACGAACTCCTCCGG and gRNA3: GCCCACACGAACTCCTCCGG. All cell lines were authenticated by one or more of short tandem repeat polymorphism, single-nucleotide polymorphism and fingerprint analyses and underwent routine mycoplasma testing by PCR.

### Cell lysis and immunoprecipitation

Cell lysates for immunoblotting and co-immunoprecipitation were prepared using radioimmuno-precipitation (RIPA) buffers and normal lysis buffers, respectively (Brummer *et al.*, 2006). Cell lysates for denatured immunoprecipitation were prepared using 1% SDS in PBS, heated at 96 °C for 10 min and then diluted to 0.1% SDS with normal lysis buffer. For immunoprecipitation, cell lysates were incubated with anti-HA affinity-agarose beads (Sigma, cat. E6679) at 4 °C overnight on a shaking platform. Following extensive washing with ice-cold lysis buffer, the immune complexes were eluted with SDS-PAGE loading buffer and subjected to Western blotting analysis.

### Western blot

Protein concentration was measured using BCA assay (Pierce, cat. 23228), and proteins subsequently separated on 8% bis‐acryl‐tris gels, and transferred to polyvinylidene difluoride membranes (Millipore, cat. IPVH00010). The membranes were blocked in 5% skim milk in TBS-Tween (TBST) (20 mM Tris pH 7.5, 150 mM NaCl, 0.05% Tween 20) for 1 h at room temperature, and then incubated with the primary antibodies at 4 °C overnight and the second antibody at room temperature for 1 h. Western blots were visualized by ChemiDoc™ Touch Gel Imaging System (Bio-Rad, cat. 1708370) and analysed by Image Lab (Version 5.2.1).

### Transfections

Plasmid transfections were performed using Lipofectamine 3000 (Life Technologies, cat. L3000015) according to the manufacturer instructions. Individual siRNAs were obtained from Dharmacon and applied to cells using DharmaFECT1 (Dharmacon, cat. T-2001-03). PEAK3 siRNAs (LQ-027257-02-0005) sequences were #18: GGACAACCCCGCUGAUCAA, #19: GGUCAGCGUCUCCAUGAUA, #20: GGCACAUCCUGGUCGCCAA, and #21: GCGGGGACGCCCUGUAUUA. PTPN12 siRNAs (LQ-008064-00-0005) sequences are #11: GGAAUUAAGUUCAGAUCUA, #12: GUAAUGGCCUGCCG AGAAU, #13: GGACACUCUUACUUGAAUU and #14: CGGGAGGUAUUCACUAUGA. Negative control siRNA was ON-TARGETplus Non-targeting pool (Dharmacon, D-001810-10-20).

### Identification of PEAK3 binding partners by MS-based proteomics

Control MCF-10A cells or cells expressing HA-tagged PEAK3 were cultured in -EGF medium (DMEM/F-12 (Gibco) supplemented with 0.5◻μg/ml hydrocortisone (Sigma), 100◻ng/ml cholera toxin (Sigma), 10◻μg/ml bovine insulin (Sigma) and 2% horse serum overnight and stimulated or not with EGF (50 ng/ml) for 5 min, and then were lysed in normal lysis buffer (Brummer *et al.*, 2006). Cell lysate (5 mg) was incubated with 60 μl of anti-HA affinity-agarose beads slurry (Sigma, cat. E6679) for 4 h at 4 °C. Following washing with lysis buffer and then wash buffer (Tris 20 mM, pH 7.4, NaCl 150 mM), the immunocomplex was digested with 1 μg trypsin (Promega, Madison, WI) in 50 mM ammonium bicarbonate solution. Tryptic digests were acidified with 10% trifluoroacetic acid (TFA) to pH 2–3, desalted with a C18 column (Thermo Fisher Scientific, Waltham, MA) and eluted with 0.1% TFA/40% acetonitrile (ACN). Peptides were dried with a SpeedVac and re-suspended in 2% ACN/0.1% formic acid (FA). The resulting peptides were analyzed by LC-MS/MS and raw files processed as previously described (Nguyen *et al*, 2019). Prism (version 8.4.2) was used to compute fold changes and raw p values using unpaired-two tailed t test. Candidate interacting proteins were defined by applying the following criteria: p value <0.05 and a fold change of >1.5 against the control IP. The mass spectrometry proteomics data have been deposited to the Proteome-Xchange Consortium via the PRIDE (Perez-Riverol *et al*, 2019) partner repository with the dataset identifier PXD023687.

### Wound healing assay

MDA-MB-231 cells were seeded in 24-well plates at 2.5 × 10^5^ cells/well in duplicate. Wounds were made by scraping with a 200 μl plastic pipette tip across the cell monolayer, and then the cells were cultured in 0.4% FBS RPMI-1640 medium with 1 μg/ml mitomycin C (Sigma, cat. M4287) to prevent cell division. Images were recorded at 0 and 24 h after scratching. Wound areas were subsequently quantified using Fiji (ImageJ) software (Version: 2.0.0).

### Three-dimensional (3D) cultures of MCF-10As

To establish 3D cultures, cells were plated into a 96-well plate (Falcon) pre-coated with 35 μl of Matrigel (Corning, cat. 354230) at a density of ~800 cells/well in medium containing 2% Matrigel (Brummer *et al.*, 2006). Cells were allowed to form acini for up to 12 days and fresh medium was replaced every 3 to 4 days. Photographs of acini were taken using a Leica DFC9000 camera with Leica Application Suite X software. Acinar size was measured using Fiji (ImageJ) software (Version: 2.0.0).

### Immunofluorescence staining of acini

Acini were fixed with 2% paraformaldehyde in PBS for 20 min and permeabilised with 0.5% Triton X-100 in PBS for 10 min at 4 °C. Acini were washed three times with PBS/glycine (130 mM NaCl, 7 mM Na2HPO4, 3.5 mM NaH2PO4, 100 mM glycine) for 15 min each. Acini were blocked for 1.5 h with primary block solution (PBS, 7.7 mM NaN3, 10% bovine serum albumin, 0.2% Triton X-100, 0.05% Tween-20). Primary antibodies were added in primary block solution overnight at 4 °C, then acini were washed three times with IF buffer (PBS, 7.7 mM NaN3, 0.1% bovine serum albumin, 0.2% Triton X-100, 0.05% Tween-20) for 20 min each with gentle rocking. Alexa Fluor® secondary antibodies were added in primary block solution for 45 min at room temperature. Acini were washed three times with IF buffer for 20 min each with rocking. Acini were washed once with PBS, then mounted onto cover slips with gold antifade reagent with DAPI (Invitrogen, cat. P3693). Images were obtained using a Nikon C1 inverted confocal microscope.

### Cell motility assay

MDA-MB-231 cells were seeded in 12-well plates at 1.5 × 10^4^ cells/well in duplicate and cultured overnight. Then the cells were cultured in 10% FBS RPMI-1640 medium with 1 μg/ml mitomycin C (Sigma, cat. M4287) to prevent cell division. Images were photographed at 8 random positions for each condition. Images of each position were taken every 20 min for 24 h using a Leica DFC9000 with Leica Application Suite X software. Cell motility was measured using Fiji (ImageJ) software (Version: 2.0.0).

### Recombinant protein expression and purification

Synthetic genes encoding full-length (FL) human Grb2 (Uniprot P62993-1; residues 1-217) and CrkII (Uniprot P46108-1; residues 1-330) were each cloned into a modified pCOLD IV vector (Takara) that contains a N-terminal His TEV cleavage tag by Genscript. Protein expression was carried out in Escherichia coli C41(DE3). Inoculated cultures were grown in Super Broth media (100 μg/mL ampicillin) at 37°C to appropriate density (OD600 ~ 0.8-1.0), then the cultures were cooled to 16 °C and protein expression induced with IPTG (0.5 mM) at 16 °C for 16-20 h. Cell pellets were harvested and stored at −80 °C prior to purification.

For purification of both Grb2-FL and CrkII-FL proteins, bacterial cells were resuspended in lysis buffer (20 mM Tris pH 7.5, 500 mM NaCl, 10% v/v glycerol, 5 mM DTT, 5 mM imidazole, 0.1% Thesit), supplemented with Complete EDTA-free Protease Inhibitor Cocktail (Roche) and lysed by sonication. The supernatant was clarified by centrifugation (45,000×g, 4 °C, 30 min), filtered and incubated in batch mode with Ni-NTA resin (Roche) (1 mL bed volume per 1L cell culture, pre-equilibrated in lysis buffer) for 2 h at 4 °C. The resin was sequentially washed with 2 × 25 mL of lysis buffer (pelleted by centrifugation at 1,500xg, 4 °C, 5 min), then resuspended in 40 mL of wash buffer (20 mM Tris pH 7.5, 500 mM NaCl, 10% (v/v) glycerol, 5 mM DTT, 5 mM imidazole) and transferred to a gravity column. After extensive washes with wash buffer, the protein was eluted in wash buffer supplemented with 250 mM imidazole, then subjected to size exclusion chromatography (SEC) (Superdex-75 16/600, GE Healthcare) in SEC buffer (20 mM Tris pH 7.5, 200 mM NaCl, 5% v/v glycerol, 0.5 mM tris (2-carboxyethyl) phosphine (TCEP)). Fractions containing monomeric forms of Grb2-FL or CrkII-FL were cleaved overnight using TEV protease then further purified by anion exchange chromatography (MonoQ 5/50 GL) (Cytiva) using Buffer A (20 mM HEPES pH 7.5, 0.5 mM TCEP) and Buffer B (20 mM HEPES pH 7.5, 1M NaCl, 0.5 mM TCEP). Fractions containing pure monomeric Grb2-FL or CrkII-FL proteins were pooled, concentrated and flash-frozen for storage at −80 °C.

### Isothermal titration calorimetry (ITC) binding experiments

ITC binding experiments were conducted using a MicroCal iTC200 instrument (Malvern Instruments). Titrations were conducted at 20 °C in standard mode (protein in cell, in syringe) and consisted of 19 injections of 2 μL peptide solution at a rate of 2 sec/μL at 90 sec time intervals, whilst stirring at 1000 rpm. An initial injection of protein (1 μL) was made and discarded during data analysis. Proteins (Grb2-FL or CrkII-FL) were first dialysed overnight at 4°C into ITC buffer (20 mM HEPES pH 7.4, 150 mM NaCl, 1 mM TCEP). Peptides were diluted into ITC buffer from 10 mM stock solutions prepared in water. Peptides (400 μM, in the syringe) were titrated into protein (20 μM or 10 μM, in the cell). Data were fitted to a single binding site model to obtain the stoichiometry (N), the dissociation constant (K_D_) and the enthalpy of binding (ΔH), using the Microcal Origin software (Version 7.0, OriginLab) provided by the manufacturer. The reported values are the mean ± S.E.M. from independent measurements (three for the pY peptide into Grb2-FL; four for the pY peptide into CrkII-FL; and two each for the negative control non-phosphorylated peptide into either protein).

## Acknowledgements

This study was supported by funding from the Australian Research Council (DP190103672) and National Health and Medical Research Council (APP1144149). We acknowledge the scientific and technical assistance of Monash Micro Imaging and Monash Biomedical Proteomics Facility, Monash University, Victoria, Australia. We would also like to thank Dr. Jason Cain from Hudson Institute of Medical Research for the A172 cell line.

## Author contributions

Conceptualization: R.J.D. Methodology: J.H., E.V.N., A.S., M.J.R., O.P., I.L., X.M. and R.J.D. Investigation: J.H., E.V.N, A.S., M.J.R., O.P. and X.M. Writing original to final draft: R.D., J.H., and X.M. with input from M.J.R, O.P., and I.L. Review and editing: R.J.D. Funding acquisition: R.J.D and I.L. Project administration and supervision: R.J.D. and X.M.

## Conflict of interest

The authors declare that they have no conflict of interest.

## Table legend

**Table EV1. Lists of PEAK3 binding partners.**

PEAK3 binding partners were identified by MS from MCF-10A_Control and MCF-10A_PEAK3 cells as described in Methods. Left list: The 109 proteins exhibiting significantly increased abundance in PEAK3 IPs versus control IPs. Right list: The 29 PEAK3 binding partners regulated by EGF treatment, with 7 up-regulated and 22 down-regulated.

**Expanded View Figure legends**

**Figure EV1. Characterization of PEAK1 KO, PEAK2 KO and DKO MCF-10A cells generated by CRISPR/Cas9 and data quantification of PEAK3 heterotypic association.**

A Western blotting was undertaken as indicated.

B Quantification of Western blots in Figure 1C (normalized to HA) and data expressed relative to WT PEAK3 which was arbitrarily set at 1. Error bars represent s.e.m., **, p<0.01, ****, p < 0.0001, #, p>0.05 by Student’s t-test. ND, not detected.

**Figure EV2. The dynamics of the PEAK3 interactome after EGF treatment.** X axis indicates EGF treatment time points (50 ng/ml). Levels of associated binding partners are normalized for the amount of HA-tagged PEAK3 in the IP and expressed as a percentage of their highest level over the time course.

**Figure EV3. A model of the full length PEAK3.** The pseudokinase domain is adopted from the crystal structure of PEAK2 (PDB 5VE6) dimer. SH2 and SH3 motifs within the PEST linker region and the pseudokinase domain are as shown. The position of a-turn (red) in PEAK1 and PEAK2 corresponds to the SH3(2) motif in PEAK3, PGAPWR. The interaction triad between Phe974 of αN1, Phe1045 of loop connecting β3-β4 (black) and Trp1382 of αL is shown in a green circle. In PEAK3, Phe974 is replaced by a leucine and is likely to maintain this interaction triad. SH3(2) mutation in PEAK3 may impact the orientation of the N-lobe with respect to the C-lobe, affecting this interaction triad and hence dimerisation. Grb2 and CrkII binding through their SH2 and SH3 domains is shown on the SH2 and SH3(1) motifs. Unresolved loops are shown as dashed lines.

**Figure EV4. Data quantification.**

A Quantification of Western blots in Figure 4B (normalized to HA). B Quantification of Western blots in Figure 4C (normalized to HA).

A and B Data expressed relative to WT PEAK3 which was arbitrarily set at 1. Error bars represent s.e.m., *, p<0.05, **, p<0.01, ****, p < 0.0001, #, p>0.05 by Student’s t-test. ND, not detected.

C. Proposed model of ASAP1 association. Grb2 bridges PEAK3 to ASAP1 by binding to the SH2 binding motif on PEAK3 and SH3 binding motif(s) on ASAP1. Other potential mechanisms contributing to ASAP1 association with PEAK3 (dotted lines) are bridging by CrkII, which binds to the SH3(1) motif and then associates with ASAP1 via SH2 or SH3 domain-mediated interactions, and direct binding via the ASAP1 SH3 domain. Note that integrity of SH3(1) is also required for efficient Y24 phosphorylation (red arrow).

**Figure EV5. Detection of endogenous PEAK3.**

A Validation of PEAK3 custom antibody. HEK293 cells were transfected with non-tagged PEAK3.

B PEAK3 was knocked out in MDA-MB-231 and RH30 cells by CRISPR/Cas9.

C PEAK3 was knocked down in A172 cells by PEAK3 siRNA pool (20 nM). Cell lysates from A–C were Western blotted as indicated.

## Notes

### Competing Interest Statement

The authors have declared no competing interest.

