## Supplemental figures for "PEAK3 pseudokinase represents a pro-migratory and -invasive signalling scaffold"

### Slide 1
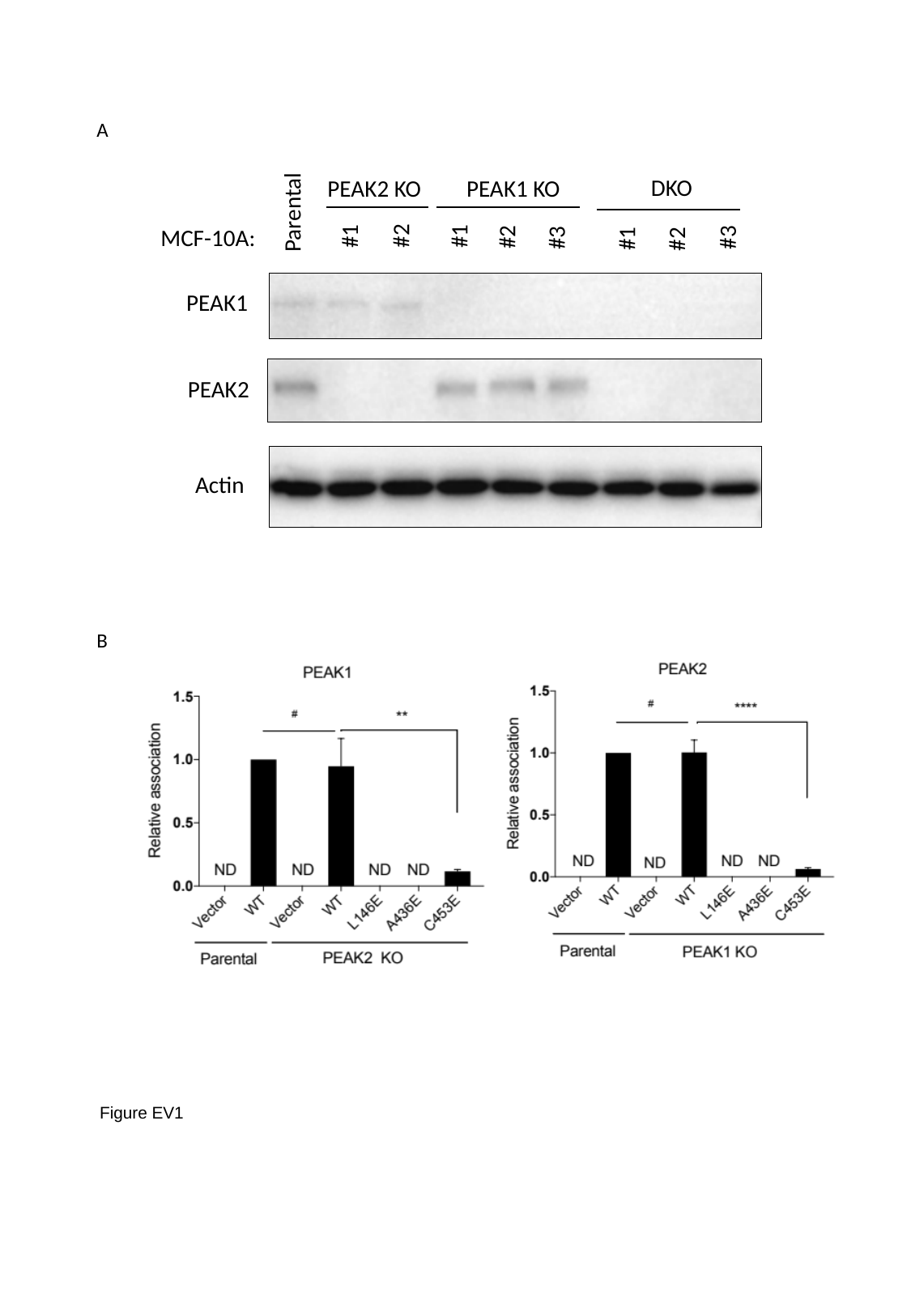

A
DKO
PEAK2 KO
PEAK1 KO
Parental
#2
#1
#1
#2
#3
MCF-10A:
#3
#1
#2
PEAK1
PEAK2
Actin
B
Figure EV1

### Slide 2
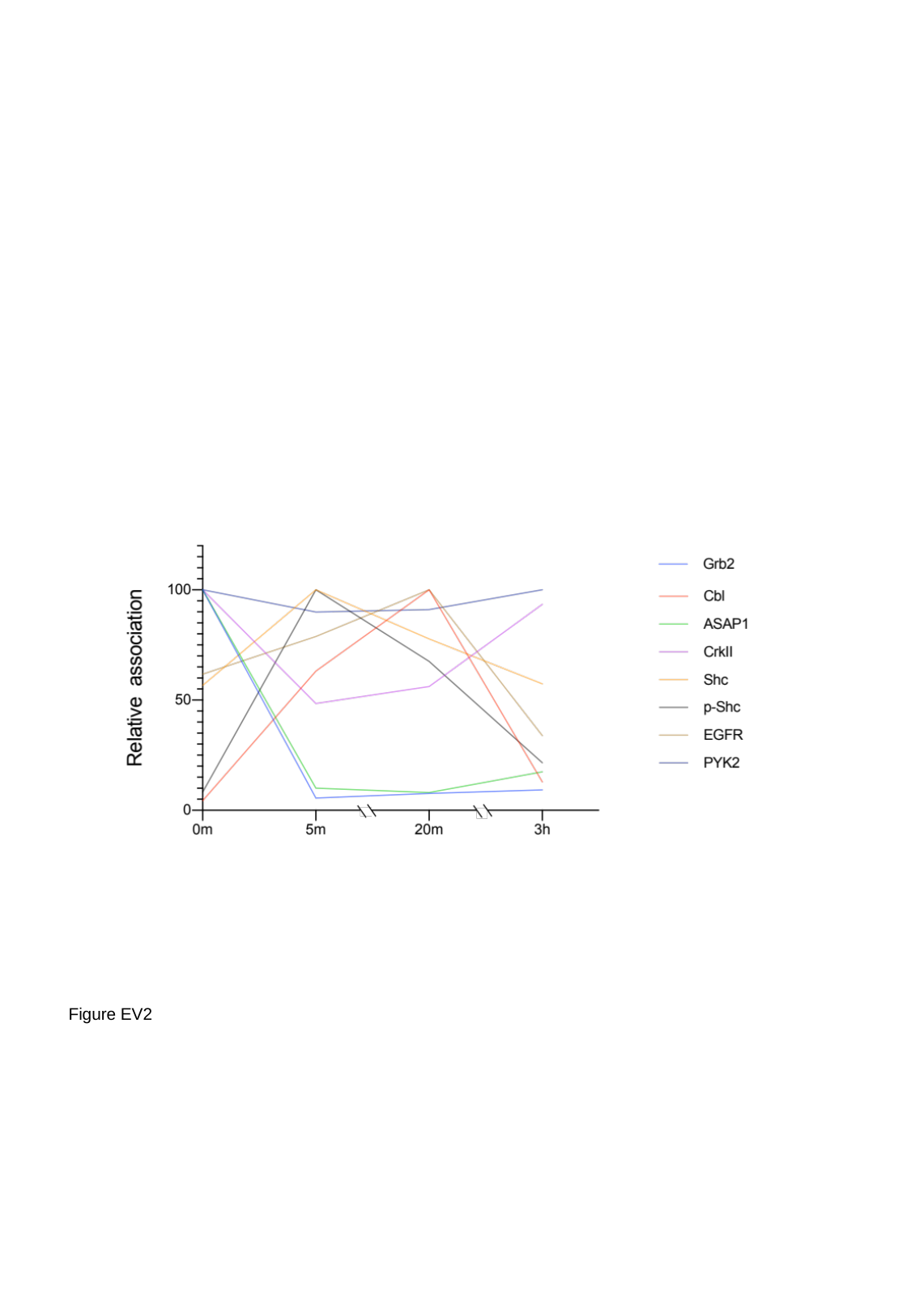

Figure EV2

### Slide 3
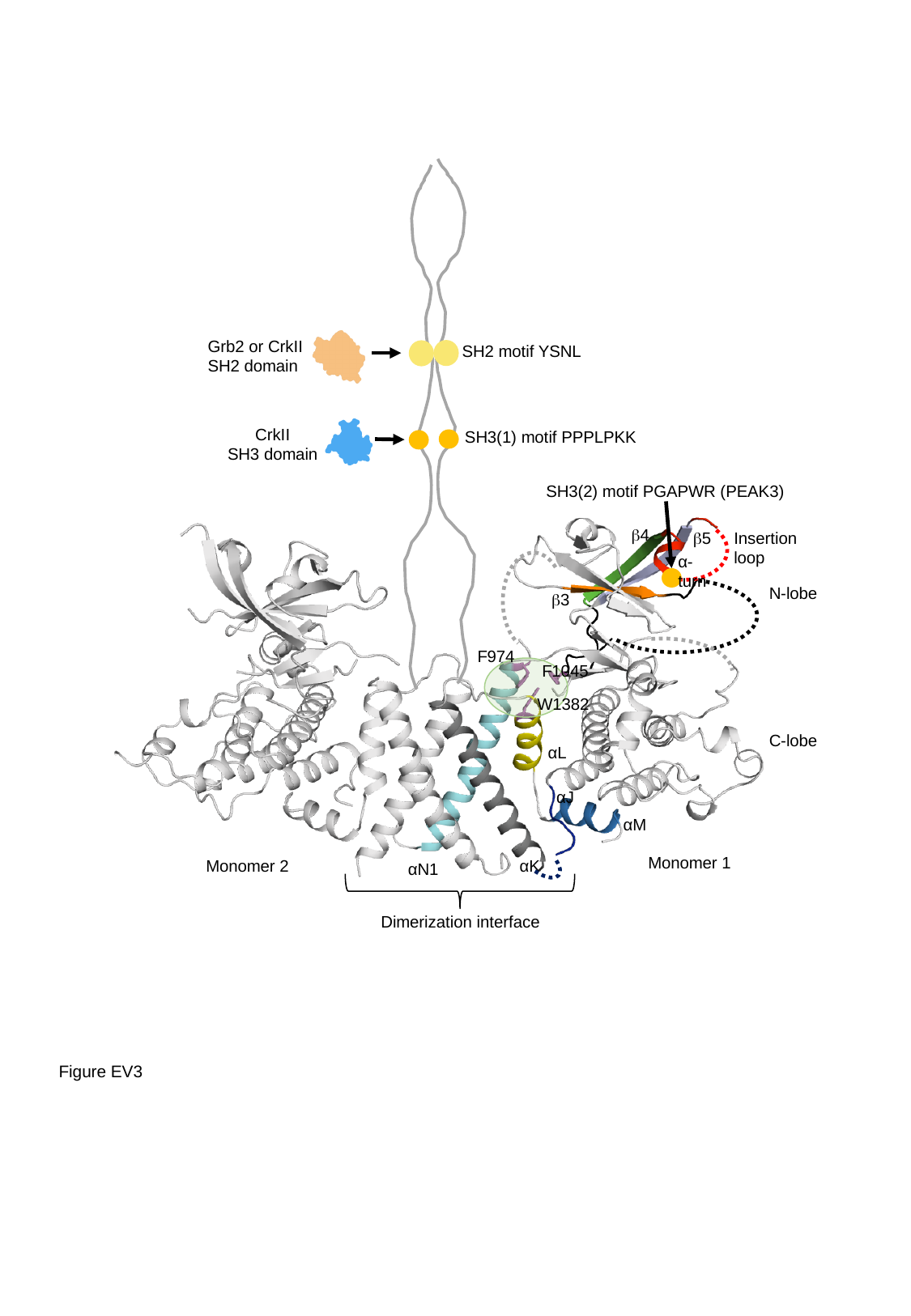

Grb2 or CrkII
SH2 domain
SH2 motif YSNL
CrkII
SH3 domain
SH3(1) motif PPPLPKK
SH3(2) motif PGAPWR (PEAK3)
β4
β5
Insertion loop
α-turn
N-lobe
β3
F974
F1045
W1382
C-lobe
αL
αJ
αM
Monomer 1
Monomer 2
αK
αN1
Dimerization interface
Figure EV3

### Slide 4
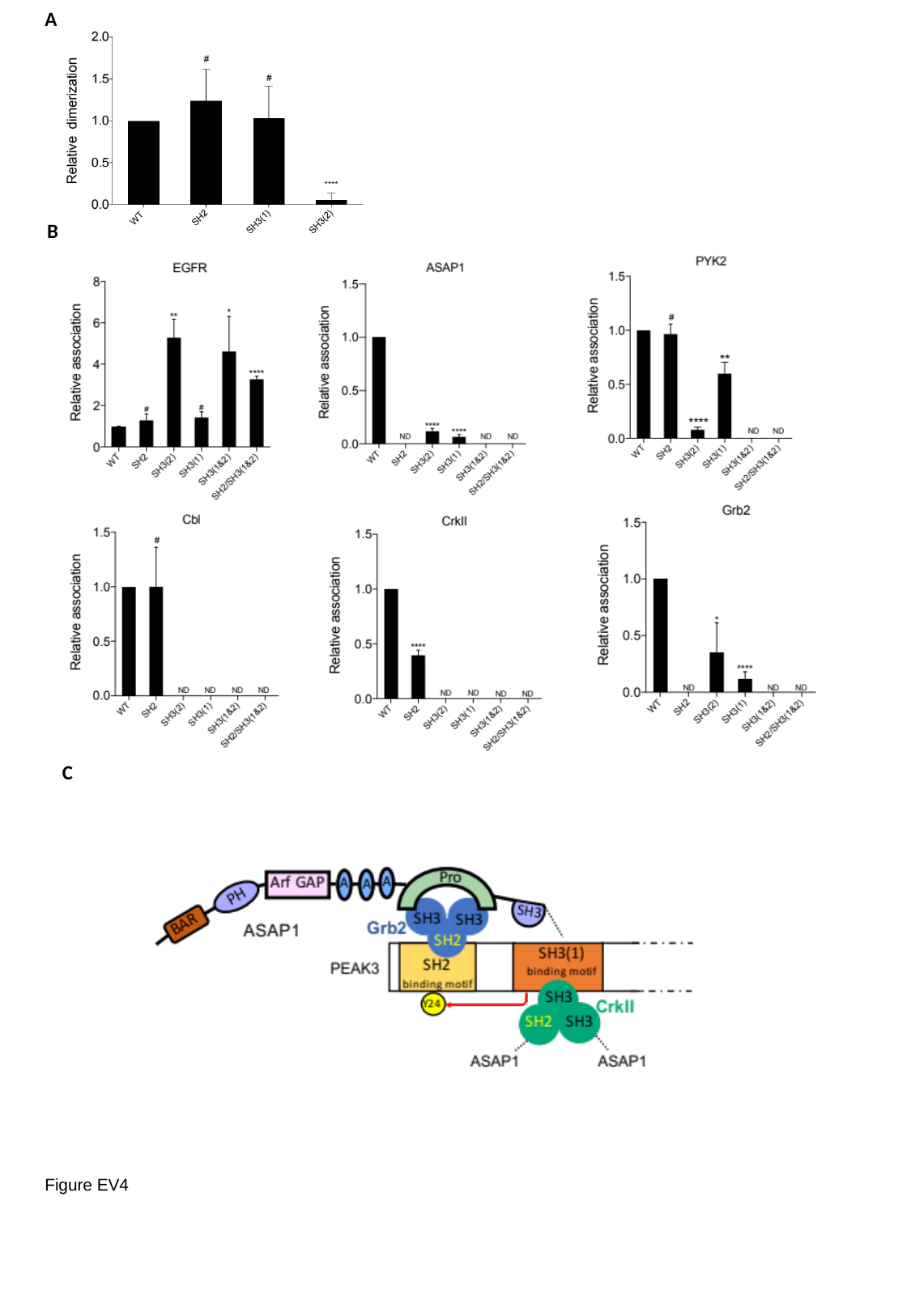

A
B
C
Figure EV4

### Slide 5
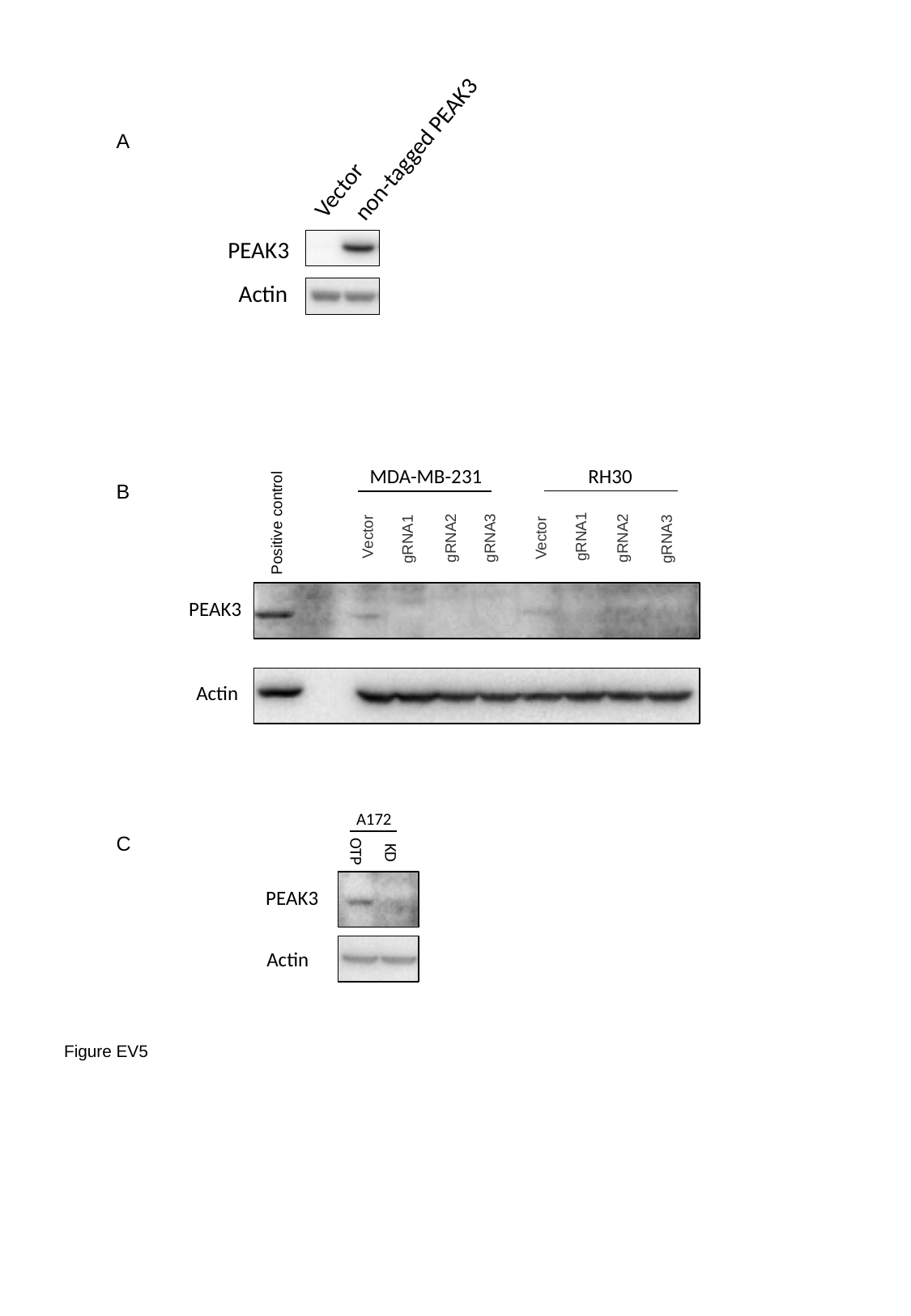

A
 non-tagged PEAK3
Vector
PEAK3
Actin
MDA-MB-231
RH30
B
Positive control
Vector
gRNA1
gRNA2
Vector
gRNA3
gRNA2
gRNA1
gRNA3
PEAK3
Actin
A172
C
OTP
KD
PEAK3
Actin
Figure EV5
